# CHD8 orchestrates chromatin landscapes during early female neuronal differentiation

**DOI:** 10.64898/2026.09.09.750428

**Authors:** Ajay Kumar Danga, Alessio Colantoni, Nicola Pomella, Nerea Ruiz Blanes, Marco Genovesi, Filiz Ersöz, Gian Gaetano Tartaglia, Morkoss M. Fakhry, Andrea Cerase

## Abstract

**Background:** Autism spectrum disorder (ASD) shows a strong male bias (∼4:1). CHD8, a frequently mutated ASD gene, regulates neuronal development and Xist regulation in X-chromosome inactivation (XCI). However, its role in female neurodevelopment remains poorly understood due to the predominance of male-derived or sex-agnostic models.

**Methods:** We performed integrated multi-omics analysis of female mouse embryonic stem cells (ES) differentiating to neuronal progenitor cells (NPCs) using wild-type, CHD8 knockdown (KD), knockout (KO), and domain-specific rescue lines (full-length, ΔChromo, ΔHelicase). RNA-seq, CHD8 and H3K4me3 ChIP-seq, and ATAC-seq datasets were integrated to assess transcriptional, chromatin-binding and -accessibility changes during differentiation.

**Results:** CHD8 occupancy was substantially remodelled during female neuronal differentiation, with 3,754 genes gaining NPCs-specific CHD8 binding predominantly at distal regulatory elements. CHD8 loss dysregulated 2,752 genes (1,134 upregulated and 1,618 downregulated), in NPCs. Differential accessibility analysis identified 4,486 chromatin regions with significant CHD8-dependent changes. Domain-specific rescue experiments showed that chromodomain and helicase activity makes distinct, non-redundant contributions to transcriptional recovery: ΔChromo-rescued only ∼ 1.0% of CHD8-dependent transcriptional changes, ΔHelicase rescued ∼ 41.1%, and full-length CHD8 rescued ∼ 70%. Integration of binding, expression, and accessibility data identified 8 high-confidence direct CHD8 target genes, and cross-referencing with the SFARI Autism Risk Gene database revealed that CHD8-dependent transcriptional changes converge on autism-related pathways.

**Conclusions:** CHD8 acts as a key regulator of female-derived neuronal differentiation, recruited to H3K4me3-marked promoters in a chromodomain-dependent manner, with helicase activity providing non-redundant regulatory capacity at a subset of targets. These findings provide a molecular framework for CHD8-dependent transcriptional regulation in female NPCs and underscore the importance of including female-derived systems in neurodevelopmental disorder research.

**Graphical Abstract:** 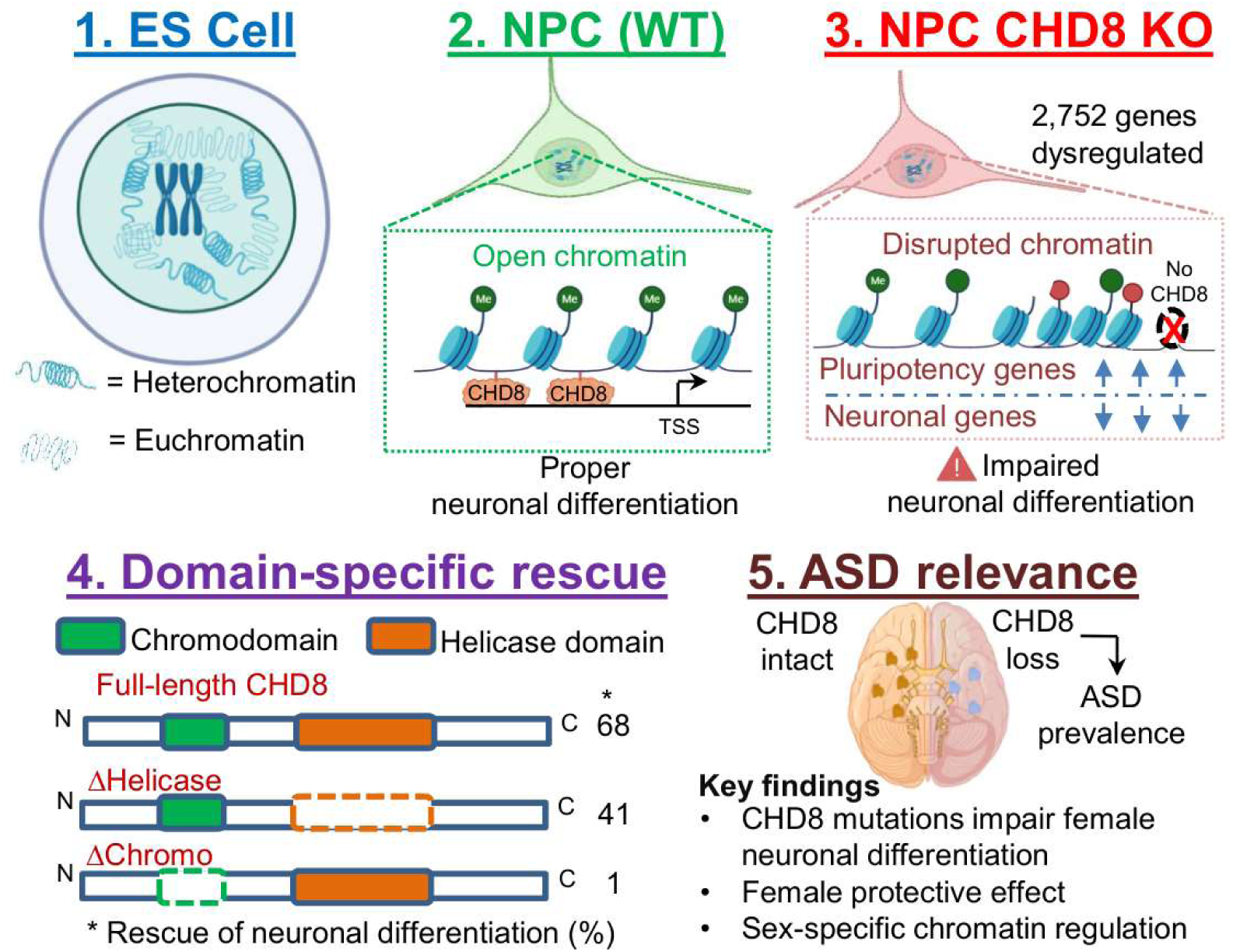

## Background

### Sex bias in autism spectrum disorder and unknown roles of Chd8 in female neurodifferentiation

Autism spectrum disorder (ASD) is a heterogeneous neurodevelopmental condition characterised by impaired social communication, restricted interests, and repetitive behaviours [1,2]. One of the most striking and poorly understood features of ASD is its pronounced male bias, with males diagnosed approximately ∼ 4 times more frequently than females [3,4]. This sex difference emerges early in development and persists across the lifespan, suggesting fundamental biological differences in neurodevelopmental trajectories between males and females [5]. Multiple hypotheses have been proposed to explain ASD’s sexual dimorphism, including the "female protective effect," differential genetic architecture, hormonal influences, and sex-specific gene expression patterns [6,7]. Intriguingly, while overall ASD diagnoses are heavily male-skewed, severe *de novo* loss-of-function mutations in chromatin remodelers like *CHD8* break through this baseline female protection, resulting in an exceptionally high phenotypic penetrance and severe clinical presentation in affected females [8]. However, the molecular mechanisms underlying these sex differences remain largely elusive. Critically, the vast majority of ASD research has been conducted using male-derived samples, male animal models, or sex-agnostic systems where biological sex is not considered as a variable [9,10]. This male-centric bias in research has resulted in a profound gap in our understanding of ASD neurobiology in females and the mechanisms generating sexual dimorphism. Chromodomain Helicase DNA-binding protein 8 (CHD8) has emerged as one of the most recurrently mutated genes in ASD genetics. *De novo* loss-of-function mutations in CHD8 are among the most frequently identified genetic causes of ASD, accounting for approximately 0.5% of cases [8,11]. Individuals with CHD8 haploinsufficiency exhibit a highly penetrant ASD phenotype, often accompanied by macrocephaly, gastrointestinal issues, and distinctive facial features [12,13]. The consistency and severity of the CHD8-ASD phenotype have made CHD8 a priority target for understanding ASD pathophysiology differences between males and females. CHD8 belongs to the CHD family of ATP-dependent chromatin remodelers and contains two N-terminal chromodomains that recognize histone H3 lysine 4 trimethylation (H3K4me3), a SNF2-like ATPase/helicase domain, and C-terminal BRK domains [14,15]. Through its chromatin remodelling activity, CHD8 regulates gene expression programs essential for brain development, including neuronal differentiation, synaptogenesis, and circuit formation [16,17]. Mouse models with heterozygous Chd8 deletion recapitulate core ASD-like behaviours, including social deficits, repetitive behaviours, and altered communication [18,19].

CHD8 research has predominantly employed male-derived cells, mixed-sex populations analysed without sex stratification, or systems where sex-specific effects cannot be assessed [20,21]. Understanding CHD8’s sex-specific roles is therefore critical for comprehending both its normal function and its contribution to ASD’s sexual dimorphism.

Female mammals possess two X chromosomes, necessitating X chromosome inactivation (XCI) to achieve dosage compensation of X-linked gene expression relative to XY males [22]. XCI is initiated early in female development [23] and in differentiating XX embryonic stem cells [24] and involves chromosome-wide transcriptional silencing of one X chromosome mediated by the long non-coding RNA Xist [25], which is transcriptionally regulated by several in cis long non-coding RNA and in trans proteins [26–40] and Xi localization at the nuclear periphery [41,42]. Importantly, XCI is not complete in humans. Approximately 15-20% of X-linked genes escape inactivation and are expressed from both X chromosomes in females [43]. These escapee genes contribute to sexual dimorphism in gene expression and may influence sex differences in brain development and disease susceptibility [44,45]. Recent evidence suggests that the dynamics and maintenance of XCI during neuronal differentiation may influence neurodevelopmental outcomes and potentially be a novel therapeutic target [46]. Disruption of XCI regulators can affect neuronal gene expression programs and differentiation trajectories [47,48]. Furthermore, many X-linked genes are involved in cognitive function and neurodevelopment, and mutations in these genes cause intellectual disability and ASD [49]. The interplay between XCI status, X-linked gene expression, and autosomal developmental programs in female neurogenesis remains poorly characterised. Beyond X-chromosome biology, female neurodevelopment differs from male neurodevelopment in numerous aspects, including timing of neurogenesis, patterns of synaptogenesis, responses to hormonal signals, and epigenetic landscapes [50,51]. Estrogen receptor signalling, for example, influences chromatin organization and gene expression in developing neurons [52]. Sex-specific transcriptional programs have been identified in the foetal human brain, with notable differences in genes related to immune function, synaptic organization, and neurotransmitter systems [53]. Despite CHD8’s prominence in ASD genetics and neurodevelopmental biology, it is still not known how CHD8 regulate female neuronal differentiation programs.

To address these questions, we performed comprehensive multi-omics profiling of female mouse ES cells undergoing neuronal differentiation under conditions of normal CHD8 expression, CHD8 reduction, CHD8 depletion, and domain-specific rescue. Our integrated analysis of transcriptomics (RNA-seq), CHD8 chromatin occupancy (ChIP-seq), active chromatin marking (H3K4me3 ChIP-seq), and chromatin accessibility (ATAC-seq) provides an integrated characterisation of CHD8’s function in female neuronal development. Here, we report that CHD8 orchestrates a complex transcriptional program essential for proper female neuronal differentiation through direct binding to thousands of promoters marked by H3K4me3. CHD8 loss causes widespread dysregulation of ASD-risk genes, synaptic function genes, and developmental regulators, with both chromodomains and helicase activity being essential for proper function. Our findings establish CHD8 as a key regulator of female neuronal differentiation and provide a framework for investigating sex-biased mechanisms in ASD.

## Methods

### Cell culture and differentiation

ES cell maintenance: Female F1 hybrid (129/Castaneus) mouse ES cells (Fa2L-S4 line) carrying a Tsix mutation were maintained in 2i conditions as previously described [54,55]. Cells were cultured on gelatin-coated plates in N2B27 medium supplemented with PD0325901 (1 μM), CHIR99021 (3 μM), and LIF (1000 U/ml).

Neuronal differentiation protocol: For differentiation to neural progenitor cells (NPCs), cells were plated at 150,000 cells per well in six-well plates pre-coated with gelatin (0.1%) and laminin (1:1000). Differentiation was induced by switching to neuronal differentiation medium (NDM) consisting of N2B27 base medium without 2i inhibitors or LIF [56]. Cells were maintained in NDM for 3 days with media changes every 24 hours. Differentiation efficiency was monitored by qRT-PCR analysis of pluripotency markers (Rex1, Sox2) and neuronal markers (Nestin).

### Generation of CHD8-deficient and rescue cell lines

CHD8 knockout line: CHD8-knockout (KO) ES cells were generated by CRISPR-Cas9 targeting as previously described [54]. Briefly, guide RNAs targeting the chromodomain and helicase-coding regions were designed and cloned into pX459 (Addgene #62988). Cells were transfected using Lipofectamine 3000, selected with puromycin (2 μg/ml) for 7-9 days, and individual clones screened by PCR and sequencing. Clone C4 carrying frameshift mutations in both domains was validated by Western blot and used for all experiments.

CHD8 knockdown lines: Stable knockdown lines were generated by lentiviral transduction with shRNA constructs (Sigma). Two independent shRNAs targeting different regions of Chd8 mRNA were used (Chd8.1: NM_201637.2-847s1c1; Chd8.2: NM_201637.2-3342s21c1), achieving ∼ 40% and ∼ 80% knockdown, respectively. Scrambled shRNA (SHC002) served as a control.

Domain-specific rescue constructs: Full-length mouse Chd8 cDNA and domain deletion mutants (ΔChromo: deletion of amino acids 1-450; ΔHelicase: deletion of amino acids 1120-1680) were cloned into the pCAG expression vector with neomycin resistance. Constructs were linearized and electroporated into CHD8-KO cells, followed by G418 selection (400 μg/ml) for 7-9 days. Rescue efficiency was verified by Western blot and expression normalized to match wild-type levels.

### RNA extraction and qRT-PCR

Total RNA was extracted using RNeasy Mini Kit (Qiagen) with on-column DNase treatment. RNA (1-2 μg) was reverse-transcribed using Maxima First Strand cDNA Synthesis Kit (Thermo Fisher). Quantitative RT-PCR was performed using KAPA SYBR FAST qPCR Master Mix on an Applied Biosystems QuantStudio system. Expression levels were normalized to GAPDH using the ^ΔΔ^Ct method. Primer sequences have been previously reported [54].

### RNA-sequencing

Library preparation: PolyA-selected RNA libraries were prepared using NEBNext Ultra II Directional RNA Library Prep Kit according to manufacturer’s instructions. Libraries were pooled and sequenced on Illumina NovaSeq 6000 platform. Post-alignment quantification yielded a mean of 35.9 million mapped read pairs per sample (ranging from 7.7 to 70.4 million mapped read pairs across samples; 80 bp paired-end reads). Bioinformatic analysis: Raw reads were pre-processed using Cutadapt v4.8 [57] with parameters *-m 50 -a AGATCGGAAGAGCACACGTCTGAACTCCAGTCA -AAGATCGGAAGAGCGTCGTGTAGGGAAAGAGTGT --nextseq-trim=20* and aligned to the mouse mm10 reference genome and Ensembl 102 transcriptome [58] using STAR v2.7.10b [59] with parameters*—quantMode TranscriptomeSAM GeneCounts --outSAMtype BAM SortedByCoordinate –readFilesCommand zcat*. Gene-level read counts were obtained from the fourth column of the *ReadsPerGene.out.tab* produced by STAR [60]. Differential expression analysis was performed using DESeq2 v1.30.0 [61]. Batch effects were controlled using the DESeq2 design formula. Fragments Per Kilobase of transcript per Million mapped reads (FPKM) values were calculated using the *fpkm* function, setting the *robust* parameter to *TRUE*. Log_2_fold-change values were shrunk using the apeglm method [62]. Genes with adjusted p-value (FDR) < 0.05 and absolute log_2_ fold-change (LFC) > 0.5 were considered differentially expressed. This threshold yielded 2,752 DEGs (1,134 upregulated, and 1,618 downregulated) for the CHD8-KO vs. WT NPCs contrast.

### ChIP-sequencing

ChIP-Seq data were generated as previously described [54].

Bioinformatic analysis: Reads were pre-processed using Trimmomatic v0.39 [63] with parameters *ILLUMINACLIP:/path/to/adapter:2:30:10:1:true SLIDINGWINDOW:20:15 MINLEN:36* and aligned to mm10 using Bowtie2 v2.3.4.1with *--very-sensitive* parameter [64]. Aligned reads were filtered using Samtools v1.1 [65] to retain only properly paired reads and exclude secondary alignments and reads with a mapping quality (MAPQ) score below 2. Duplicate reads were removed using Picard v2.25.1 MarkDuplicates tool (available at http://broadinstitute.github.io/picard), with parameters *OPTICAL_DUPLICATE_PIXEL_DISTANCE=2500 VALIDATION_STRINGENCY=SILENT MAX_SEQUENCES_FOR_DISK_READ_ENDS_MAP=50000 MAX_FILE_HANDLES_FOR_READ_ENDS_MAP=8000 SORTING_COLLECTION_SIZE_RATIO=0.25 REMOVE_DUPLICATES=true*. BEDTools [66] v2.30.0 pairToBed tool was used to remove reads mapping to ENCODE Blacklist regions [67]. Peak calling was performed using MACS2 v2.2.7 callpeak [68] with parameters*-f BAMPE --call-summits -g mm --keep-dup all* and using the corresponding input samples as controls. Peaks overlapping with IgG control peaks were removed from the IP peak sets using BEDTools intersect.Consensus peak sets across biological replicates were generated using BEDTools intersect [69,70], requiring a peak to be present in at least 2 of the biological replicates contributing to each condition; reads were then counted over this consensus peak set using BEDTools multicov. Peaks were annotated to genomic features using ChIPseeker v1.30.0 [71]. Differential binding analysis was performed directly on the BEDTools-derived count matrix using DESeq2, with regions at FDR < 0.05 considered differentially accessible. Coverage tracks were generated for visualization purposes using deepTools bamCoverage.

For all downstream analyses, peaks were restricted to standard chromosomes (chr1 – chr19, chrX, chrY), removing scaffold contigs. Peak overlap between ES and NPC conditions was computed using BEDTools intersect [66]; a peak was classified as shared if it overlapped any peak in the other condition by ≥ 1 bp (NPC perspective). This yielded 32,017 ES-cell peaks and 48,735 NPC peaks on standard chromosomes, with 10,418 (32.5%) ES-specific, 28,690 (58.9%) NPC-specific, and 20,045 shared (NPC perspective) **Supplementary Table: S1**.

### ATAC-sequencing

ATAC-seq data were generated as previously described [54].

Bioinformatics analysis: ATAC-seq reads were processed similarly to ChIP-seq reads (see above). Aligned reads were additionally shifted as described in [72] using the deepTools alignmentSieve tool with the –*ATACshift* parameter. Peaks were called using MACS2 with parameters *-f BAMPE --call-summits -g mm --keep-dup all* and no control sample. Principal component analysis of read counts across all six samples was performed on DESeq2-normalized counts. PC1 and PC2 explained 58% and 38% of total variance, respectively. One KO replicate (KO1, replicate 1) was identified as a clear outlier, positioned distant from all other samples along PC1 (and separated from its own paired replicate, KO1_rep2, by roughly 11 units on PC1 and 10 units on PC2) while wild-type and KO2 replicate pairs each clustered tightly together; KO1_rep1 was therefore excluded from all downstream analyses.

A consensus peak set was constructed using DiffBind (v3.4.0) from four samples — two WT replicates and one replicate from each of the two independent CHD8-KO clones — requiring peaks to be present in at least 2 of the 4 samples, irrespective of condition (minOverlap = 2), yielding 74,486 consensus regions. Read counts at these regions were then quantified for all five non-outlier samples (two WT replicates and three KO replicates, including the second replicate of the excluded clone) using first-in-pair, quality-filtered fragments (samtools view -f 0×40 -F 0×904) via bedtools multicov. Differential accessibility analysis was performed using DESeq2 (v1.30.0), testing CHD8-KO versus WT. All retained ATAC-seq samples showed high signal-to-noise quality: FRiP scores ranged from 0.297 to 0.485, exceeding the ENCODE-recommended minimum of 0.20, and 10.1 – 10.7% of reads mapped within ± 2 kb of annotated TSS across samples, consistent with well-enriched ATAC-seq libraries. Of 74,486 consensus regions, 4,486 showed significant CHD8-dependent accessibility changes (adjusted P < 0.05). For integrative analyses linking accessibility changes to nearby genes, regions were additionally filtered to |log_2_FC| > 0.5, yielding 3,161 KO-lost and 1,325 KO-gained peaks. Genes within ± 2 kb of TSS were associated with ATAC-seq peaks using pybedtools intersect [73].

### Gene ontology and pathway enrichment analysis

Gene Ontology (GO) enrichment analysis was performed using the Enrichr API via gseapy [74] with the GO_Biological_Process_2023 gene set database. For NPC-specific target enrichment, the input gene list comprised 254 genes at the intersection of NPC-specific CHD8 ChIP-seq targets and CHD8-KO DEGs (adjusted P < 0.05). GO enrichment of significantly downregulated in CHD8-KO NPCs (adjusted P < 0.05, log_2_FC < −0.5; n = 1,618) is provided in **Supplementary Table: S2**. For both analyses, a background of all express d genes with a baseMean >10 (n = 13,590) was utilized. Because Benjamini–Hochberg FDR correction was overly conservative at these specific gene list sizes, nominal P < 0.05 was used as the significance threshold.

### Integration of multi-omics datasets

#### NPC-specific CHD8 target identification

NPC-specific CHD8 target genes were defined as genes with a CHD8 ChIP-seq peak within ± 2 kb of their annotated TSS (mm10 RefGene) exclusively in NPCs (absent in ES cells), identified using pybedtools subtract and intersect. This yielded 3,754 NPC-specific target genes. A subset of 254 genes at the intersection with CHD8-KO DEGs (adjusted P < 0.05, |log_2_FC| > 0.5, baseMean> 10) was used as input for GO enrichment analyses.

#### Direct CHD8 target classification

Genes were classified as direct CHD8 targets if they met the following two criteria: (1) NPC-specific CHD8 ChIP-seq peak within ± 2 kb of TSS (n = 3,754 gene targets); (2) differentially expressed in CHD8-KO versus WT NPCs (DESeq2 adjusted P < 0.05, |log_2_FC| > 0.5, baseMean > 10). This two-criterion intersection yielded 254 direct CHD8 target genes. A high-confidence subset of 8 genes additionally showed significant differential chromatin accessibility at proximal ATAC-seq regions (DiffBind/DESeq2, adjusted P < 0.05, |log_2_FC| > 0.5, on a balanced four-sample consensus peak set with one outlier replicate excluded — see ATAC-seq differential accessibility methods) and is used for triple-overlap analyses in **Supplementary Figure 1**.

**Supplementary Figure 1.**
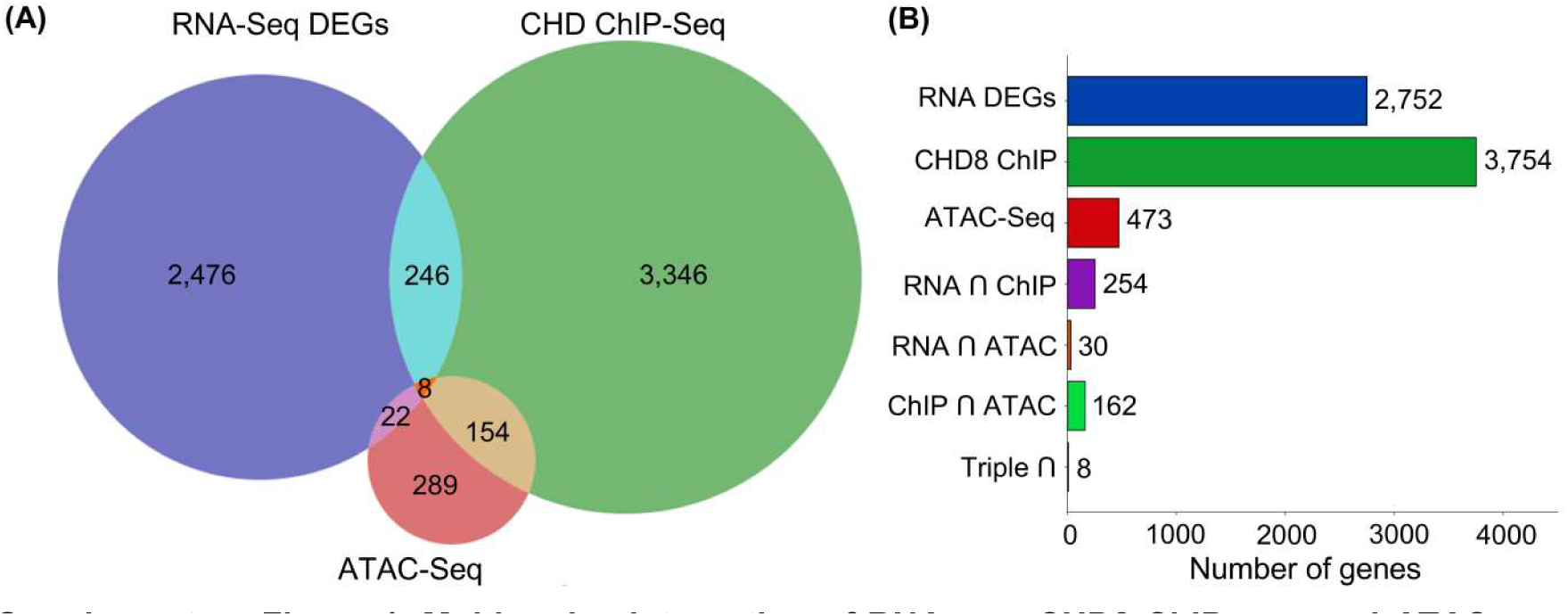
Multi-omics integration of RNA-seq, CHD8 ChIP-seq, and ATAC-seq. **(A)** Three-way Venn diagram showing overlap between RNA-seq differentially expressed genes (DEGs; adjusted *P* < 0.05, |log2FC| > 0.5; n = 2,752), CHD8 NPC-specific ChIP-seq targets (n = 3,754; peak within ± 2 kb of TSS, NPC-specific peaks only), and genes associated with significant ATAC-seq accessibility changes in CHD8-KO NPCs (DiffBind/DESeq2, adjusted P < 0.05, |log_2_FC| > 0.5; n = 473). Intersections yielded 8 genes common to all three datasets, representing high-confidence direct CHD8 targets that are simultaneously CHD8-bound, transcriptionally dysregulated, and linked to chromatin accessibility changes in KO cells. Remaining overlaps are partitioned as RNA-only (2,476), ChIP-only (3,346), ATAC-only (289), RNA ∩ ChIP (246), RNA ∩ ATAC (22), and ChIP ∩ ATAC (154). **(B)** Bar plot summarizing the size of each set and pairwise intersection from panel (A), highlighting the relative contributions of and degree of overlap between transcriptomic, binding, and chromatin accessibility datasets. TSS annotations were derived from UCSC mm10 RefGene; all intersections were computed using pybedtools.

#### Rescue fraction analysis

For each gene with |log_2_FC_KO| > 0.5, rescue fraction (RF) was calculated as: RF = (log_2_FC_rescue − log_2_FC_KO) / (0 − log_2_FC_KO) where RF = 1.0 indicates complete restoration to WT levels and RF = 0.0 indicates no rescue. RF values were clipped to the range [−1, 2] to limit the influence of outliers. A gene was classified as rescued if RF ≥ 0.5. Rescue analyses were performed for full-length CHD8 (FL), ΔChromo, and ΔHelicase constructs across 3,754 NPC-specific CHD8 target genes with |log_2_FC_KO| > 0.5. Statistical comparisons of RF distributions between constructs were performed using the Friedman test (omnibus; scipy.stats.friedmanchisquare) followed by pairwise Wilcoxon signed-rank tests (scipy.stats.wilcoxon; alternative = ‘two-sided’) with Benjamini–Hochberg FDR correction (statsmodels.multipletests v0.14.4). Categorical rescue classification used RF ≥ 0.5 per construct; non-uniform distribution across categories was assessed by chi-square goodness-of-fit test (scipy.stats.chisquare).

### Statistical analysis

All statistical tests were two-tailed unless otherwise noted. For RNA-seq and ChIP-seq, multiple testing corrections were applied using Benjamini-Hochberg method. For qRT-PCR and quantitative IF, unpaired t-tests or one-way ANOVA with post-hoc Tukey correction were used as appropriate. Sample sizes and statistical tests are indicated in figure legends. Genes with baseMean ≤ 10 were excluded from all DESeq2 analyses to remove unreliably quantified transcripts. For chromatin accessibility analysis, differential accessibility was assessed using DiffBind (consensus peak set, minOverlap = 2 across a balanced four-sample design) and DESeq2, following exclusion of one replicate identified as a technical outlier by PCA. For rescue fraction comparisons, the Friedman test and pairwise Wilcoxon signed-rank tests with Benjamini–Hochberg correction were used (scipy 1.16.3). For enrichment analysis, Fisher’s exact test was used for categorical overlap comparisons (scipy.stats.fisher_exact). Mann–Whitney U tests (two-sided) were used for non-parametric comparisons of continuous distributions. Chi-square goodness-of-fit tests were used for categorical rescue classification. Sample sizes and specific statistical tests are indicated in figure legends. Data visualization and statistical analyses were performed in Python v3.12 using pandas v2.2.2, numpy v2.0.2, scipy v1.16.3, seaborn v.13.2, matplotlib v3.10.0, pybedtools v0.12.0. Multi-omics overlaps were visualized using matplotlib-venn v1.1.2. Additional heatmap visualization and specialized genomics plots were generated in R v4.1.2 using ggplot2, ComplexHeatmap, custom scripts and TF Enrichment Analysis by (GSEAPY/ChEA). Gene identifier conversion from ENSEMBL IDs to official Gene Symbols was performed using the mygene (v3.2.2) Python client to access the BioThings API (biothings.client).

## Results

### CHD8 undergoes dynamic redistribution during female ES-to-NPC differentiation

To characterize CHD8’s role in female neuronal differentiation, we first mapped CHD8 chromatin occupancy in undifferentiated female ES cells and after 3 days of neuronal differentiation to the NPC stage [54]. In undifferentiated ES cells, we identified 32,017 high-confidence CHD8 peaks, while differentiated NPCs showed 48,735 CHD8 peaks (**Supplementary Table: S1**). This expansion of CHD8 occupancy reflects active redistribution linked to protein increase during NPCs differentiation [54]. Consequently, the simultaneous loss of 10,418 ES-specific peaks alongside gain of 28,690 NPC-specific peaks is biologically inconsistent with a simple data-scaling effect. Comparison of ES and NPC CHD8 binding patterns revealed three categories: (1) ES-specific peaks (32.5 % of ES total), (2) NPC-specific peaks (58.9% of NPC total), and (3) shared peaks present in both states (20,045 NPC peaks sharing a standard core genomic intersection of ≥ 1 bp with any ES peak; 21,599 ES peaks also overlap NPC peaks from the ES perspective — asymmetry of 1,554 reflects larger average ES peak width) (**Figure 1A**).

**Figure 1.**
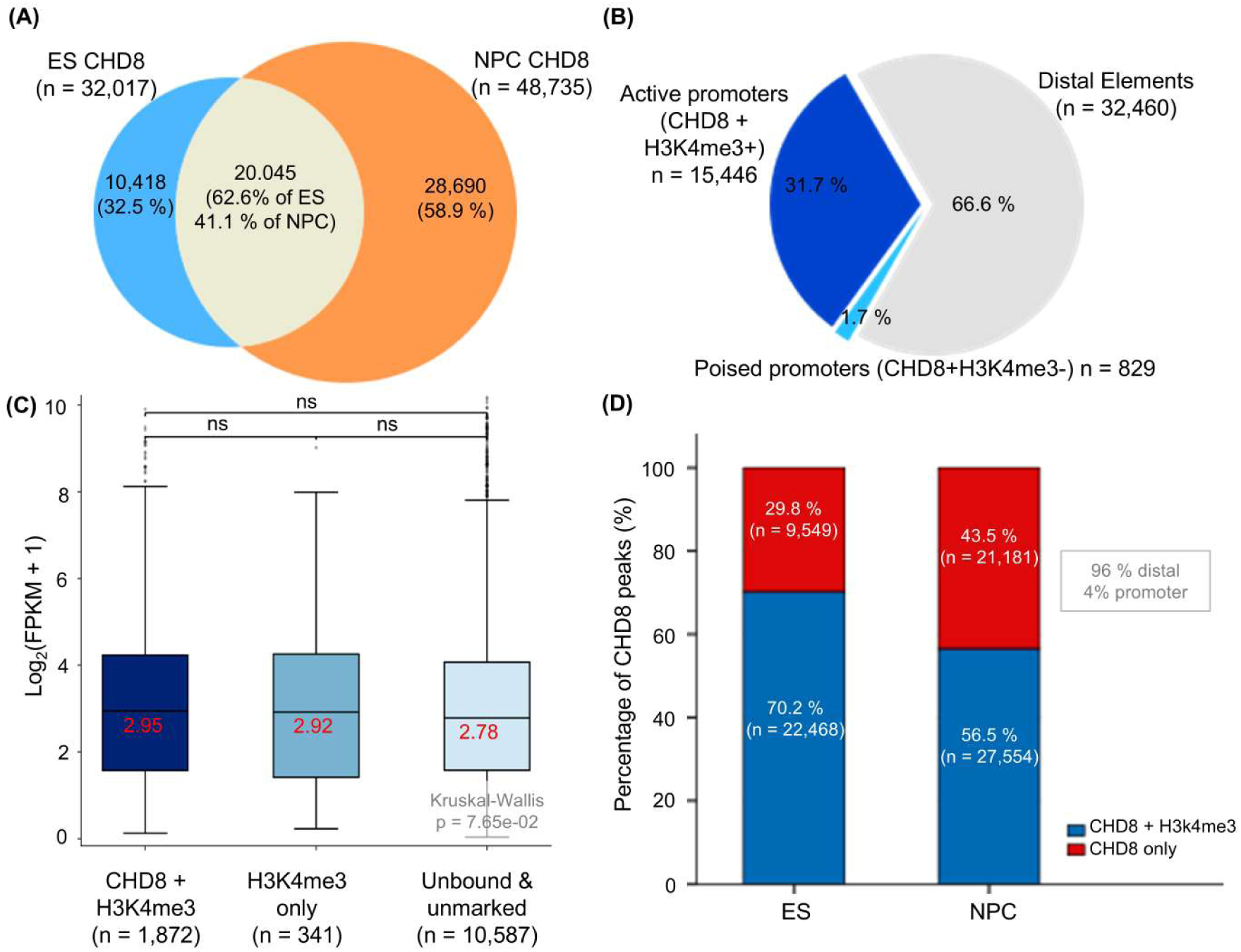
CHD8 binding dynamics during neuronal differentiation. **(A)** Venn diagram showing the overlap of CHD8 ChIP-seq peaks between embryonic stem cells (ESCs; n = 32,017) and neural progenitor cells (NPCs; n = 48,735). A substantial fraction of binding sites is cell-type specific, with 10,418 ESC-specific peaks (32.5%) and 28,690 NPC-specific peaks (58.9%). The higher number of NPC-specific sites indicates extensive remodelling of the CHD8 cistrome during neuronal differentiation. **(B)** Genomic distribution of CHD8 binding sites in NPCs. The majority of peaks localise to distal regulatory elements (66.6%), with a substantial fraction at active promoters marked by H3K4me3 (31.7%) and a small proportion at poised promoters (1.7%). These results indicate that CHD8 predominantly occupies distal regulatory regions while also associating with actively transcribed promoters. **(C)** Box-plots showing the distribution of gene expression levels (log_2_(FPKM+1), WT NPC) for genes associated with three classes of genomic regions: co-occupied by CHD8 and H3K4me3 (dark blue; n = 1,872), H3K4me3-only (light blue; n = 341), and unbound & unmarked regions (light gray; n = 10,587). Boxes indicate median and inter-quartile range. Statistical comparisons between groups are shown above the plots (ns, not significant; MWU FDR values indicated). No pairwise comparison reached statistical significance (CHD8+H3K4me3 vs. unbound & unmarked, FDR = 8.03×10⁻²; CHD8+H3K4me3 vs. H3K4me3-only, FDR = 0.76; H3K4me3-only vs. unbound & unmarked, FDR = 0.76), indicating that CHD8/H3K4me3 promoter co-occupancy is not associated with a detectable difference in gene expression. Genes were filtered to FPKM > 0.5 in at least one condition prior to analysis. Kruskal-Wallis p = 7.65×10⁻². **(D)** Stacked bar plots showing the genome distribution of CHD8 peaks based on H3K4me3 co-occupancy in embryonic stem (ES) cells and neural progenitor cells (NPCs). Blue indicates CHD8 peaks overlapping H3K4me3, and red indicates CHD8-only peaks lacking H3K4me3 signal. Percentages and counts are indicated within each bar. In ES cells, 70.2% (n = 22,468) of CHD8 peaks co-occur with H3K4me3, compared to 56.5% (n = 27,554) in NPCs. Conversely, CHD8-only peaks are more prevalent in NPCs (43.5%; n = 21,181) than in ES cells (29.8%; n = 9,549), with most CHD8-only peaks located in distal regions (∼ 96%) rather than promoters (∼ 4%).

Genomic annotation of NPC CHD8 peaks showed that CHD8 occupancy spans active promoters, poised promoters, and distal regulatory elements, classified based on H3K4me3 ChIP-seq peak co-occupancy within ± 2 kb of the TSS: peaks were designated as active promoters if a CHD8 peak overlapped both a TSS (± 2 kb)and an H3K4me3 ChIP-seq peak (CHD8+H3K4me3+; 31.7%), poised promoters lacking H3K4me3 (CHD8+H3K4me3−; 1.7%), and as distal regulatory elements if outside TSS-proximal regions (66.6%) (**Figure1B**). Approximately one-third of CHD8 NPC occupancy is therefore associated with potentially-transcribed H3K4me3-marked promoters, consistent with CHD8’s established role in chromodomain-H3K4me3-dependent recruitment. We next examined the relationship between CHD8/H3K4me3 co-occupancy and gene expression in WT NPCs, assessed using average FPKM values (FPKM > 0.5 in at least one condition) from our RNA-seq differential expression analysis (described in the following section). Genes bound by both CHD8 and enriched for H3K4me3 at their promoters did not show a statistically significant difference in expression relative to unbound and unmarked genes (median log_2_(FPKM+1) CHD8+H3K4me3 = 2.949, n = 1,872; unbound and unmarked = 2.784, n = 10,587; MWU FDR = 8.03×10^-2^; Kruskal-Wallis p = 7.65×10^-2^ (**Figure 1C**). Together, these results indicate that CHD8/H3K4me3 promoter co-occupancy is not associated with a detectable difference in steady-state expression level in WT NPCs, suggesting that CHD8 promoter binding does not, on its own, predict transcriptional output at this level of resolution.

Interestingly, we identified a subset of CHD8 peaks lacking detectable H3K4me3 co-occupancy in both ES cells (29.8%; n = 9,549 of 32,017 peaks) and NPCs (43.5%; n = 21,181 of 48,735 peaks) (**Figure 1D**). The higher proportion of CHD8-only peaks in NPCs likely reflects two converging factors. First, there is a substantial gain of NPC-specific CHD8 binding at distal regulatory regions (28,690 NPC-specific peaks; 58.9% of NPC total). Second, the overall number of H3K4me3 peaks is markedly lower in NPCs compared to ES cells (36,167 vs 84,278), reducing the likelihood of CHD8/H3K4me3 co-occupancy in the NPC state. Together, these changes indicate a redistribution of CHD8 binding from promoter-associated regions toward distal regulatory loci during neuronal differentiation. Notably, a large fraction of these CHD8-only peaks corresponds to NPC-specific gained sites, further supporting the expansion of CHD8 occupancy at distal elements. Consistent with this, CHD8-only peaks were predominantly located at distal regulatory regions (96.1% outside TSS ± 2 kb windows), with only 3.9% overlapping annotated promoters, suggesting a potential role in enhancer-mediated regulation, possibly in the absence of H3K4me3 mark.

### CHD8 loss causes widespread transcriptional dysregulation in female NPCs

To assess CHD8’s functional requirement for proper gene regulation during female neuronal differentiation, we analyzed RNA-seq data from wild-type (WT) ES cells, WT NPCs, CHD8-KO ES cells, CHD8-KO NPCs, and two independent CHD8-KD lines (Chd8.1 ∼ 40% KD; Chd8.2 ∼ 80% KD) at the NPC stage from previously-published work [54] (**Supplementary Figure 2**). Differential expression analysis (DESeq2; |log2FC| > 0.5, adjusted P < 0.05; 13,590 genes retained after filtering for an average normalized read count across samples (baseMean > 10) revealed substantial transcriptional dysregulation across all CHD8 perturbation conditions. CHD8-KO NPCs showed 1,134 upregulated and 1,618 downregulated genes compared to WT NPCs (total 2,752 DEGs; **Supplementary Table: S3**). KD line 1 (8.1 KD) exhibited 43 upregulated and 15 downregulated genes (total 58 DEGs; **Supplementary Table: S4**), and KD line 2 (8.2KD) showed 188 upregulated and 233 downregulated genes compared to scrambled shRNA controls (total 421 DEGs; **Figure 2**).

**Figure 2.**
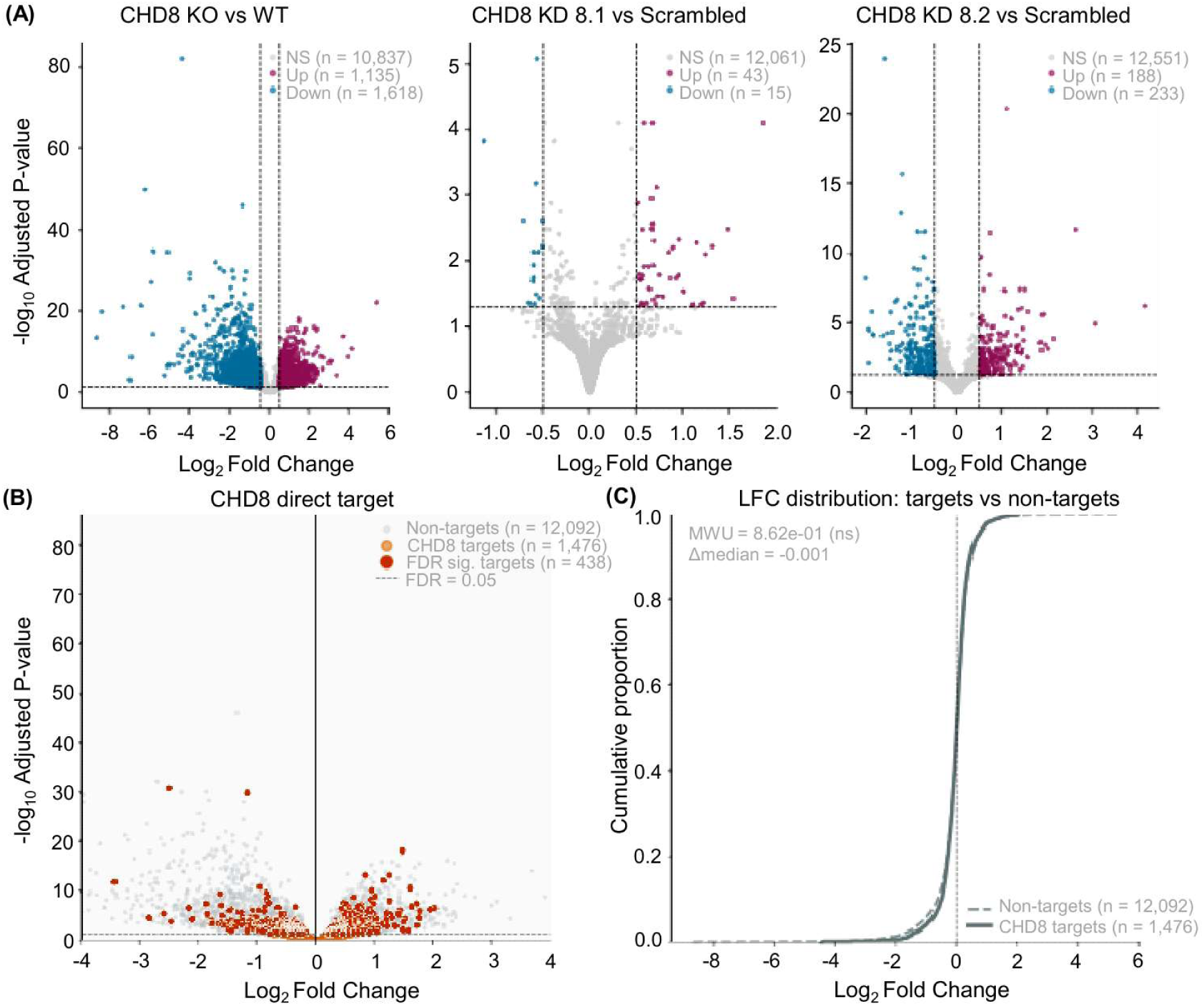
Transcriptional consequences of CHD8 loss in neural progenitor cells. **(A)** Volcano plots showing differential gene expression across independent CHD8 perturbation models: knockout (KO vs WT), and two shRNA-mediated knockdowns (KD 8.1 and KD 8.2 vs scrambled controls). Genes are coloured by direction of change (upregulated, downregulated, or non-significant). Consistent patterns of differential expression across models support the reproducibility of CHD8-dependent transcriptional effects. **(B)** Volcano plot showing differential gene expression in CHD8 knockout (KO) versus wild-type (WT) NPCs. CHD8 direct target genes (defined by NPC-specific CHD8 binding within 2 kb of the transcription start site) are highlighted in orange, with significantly dysregulated targets (FDR < 0.05) shown in darker orange. CHD8-bound genes are distributed across both up- and downregulated regions, indicating heterogeneous transcriptional responses. **(C)** Cumulative distribution of log_2_ fold-change values for CHD8 direct target genes compared to non-target genes. No significant difference between the distributions was observed (Mann–Whitney U test, p > 0.05), indicating that CHD8 binding alone does not confer a uniform directional effect on gene expression.

Having established the global transcriptional landscape in Chd8 depleted cells, we next asked whether CHD8’s transcriptional impact is preferentially concentrated at its direct genomic targets. CHD8 direct NPC target genes defined by NPC-specific ChIP-seq peaks within 2 kb of the TSS (n = 1,476 detected in RNA-seq after baseMean > 10 filtering, of which 438 reach FDR < 0.05) are highlighted within the KO vs WT volcano plot (**Figure 2B**). These direct targets are distributed across the full range of fold-change and significance values, rather than clustering among the most strongly dysregulated genes, reflecting the complex and context-dependent transcriptional role of CHD8 at NPC loci.

A two-sided Mann-Whitney U test comparing the log_2_ fold-change distributions of direct NPC target genes (n = 1,476 detected in RNA-seq) against non-target genes (n = 12,092) revealed no global shift between the two classes (p = 0.862, Δmedian log_2_FC = −0.001), demonstrating that physical promoter occupancy by CHD8 does not inherently dictate a uniform directional bias on baseline transcriptional output. This is consistent with our earlier finding that CHD8 ChIP-seq-derived binding does not, by itself, predict gene deregulation, suggesting that CHD8’s transcriptional influence is highly context-dependent, or linked to secondary events and cannot be captured by binding alone [17,75] **(Figure 2C)**.

The lower DEG count in CHD8.1^KD^ relative to CHD8.2 is consistent with its less efficient CHD8 depletion, as confirmed by qRT-PCR and western blot (**Supplementary Figure 2**), and is consistent with the dose-dependent relationship between CHD8 levels and transcriptional dysregulation previously reported [54].

**Supplementary Figure 2.**
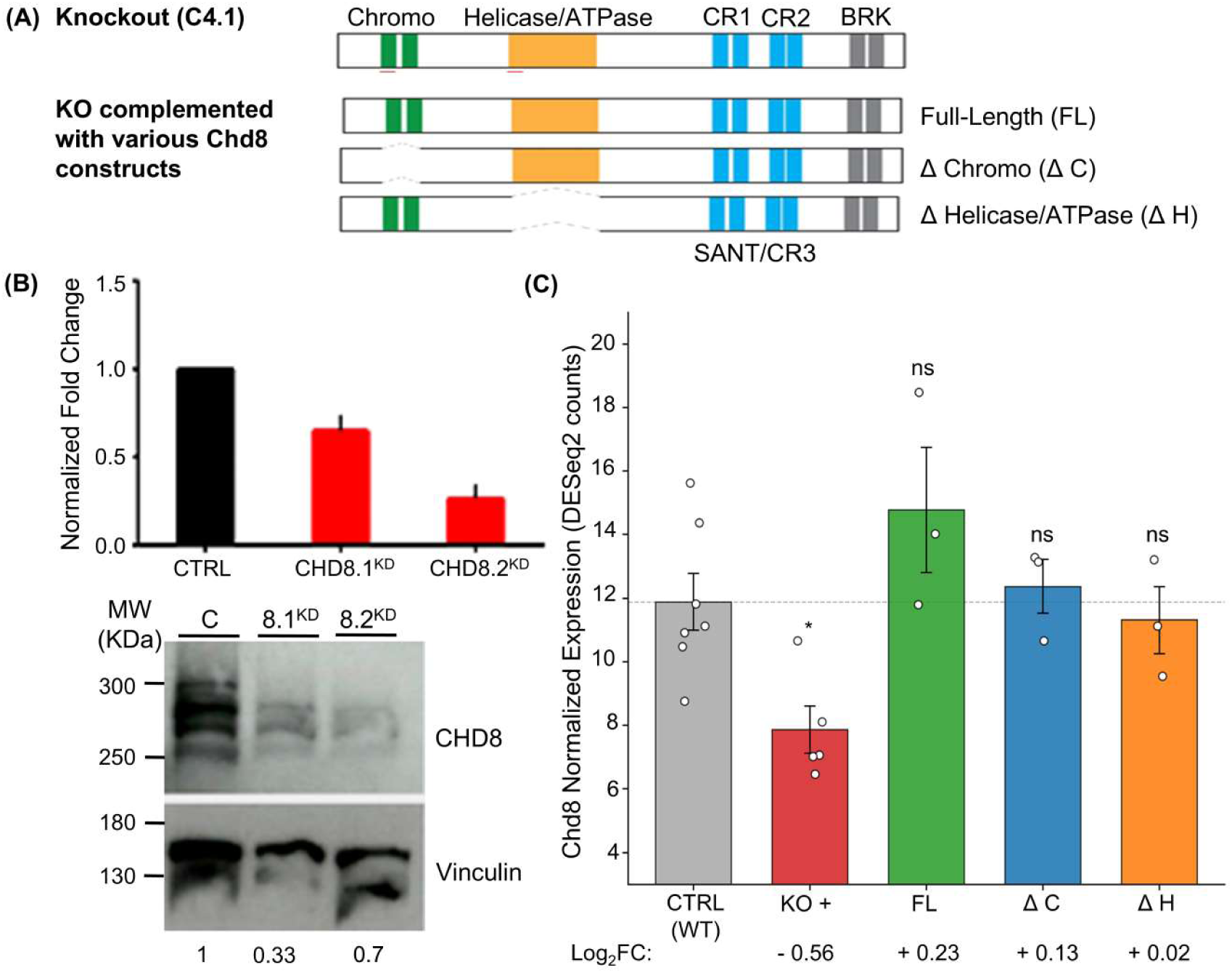
Characterization of CHD8 Mutant and Rescue Cell Lines. **(A)** Schematic representation of CHD8 protein domains and the specific deletions for ΔChromo (ΔC) and ΔHelicase/ATPase (ΔH) rescue constructs. Red lines under the domains represent the position of the guides used to generate the CHD8 KO cells. Validation of shRNA-mediated knockdown (8.1^KD^ and 8.2^KD^) via qRT-PCR **(B)** and Western blot showing a dose-dependent reduction in CHD8 levels. [Adapted from Cerase et al., Communications Biology, 2021 (DOI: 10.1038/s42003-021-01945-1), under Creative Commons Attribution 4.0 International License (CC BY 4.0)]. **(C)** DESeq2-normalised CHD8 expression in wild-type CTRL (grey), CHD8 knockout (KO, red), full-length rescue (FL, green), chromodomain-deletion rescue (ΔC, blue), and helicase-domain-deletion rescue (ΔH, orange) NPCs. Bars = mean ± SEM; dots = individual biologicalreplicates (CTRL n = 7; KO n = 5; FL/ΔC/ΔH n = 3). Significance vs CTRL by DESeq2 Wald test with Benjamini–Hochberg correction: *p < 0.05, ns = not significant.

To confirm that the observed transcriptional changes reflect specific CHD8 loss rather than an artifact of a single technical approach; we assessed the global concordance of log_2_ fold-change values across our primary depletion models. Pairwise comparison between the complete CRISPR-mediated homozygous knockout (KO) and the intermediate knockdown model (KD clone 8.1) revealed a weak but significant positive correlation (r = 0.19, P < 0.001), consistent with the comparatively larger dynamic range and milder depletion efficiency expected of this clone. A somewhat stronger, through still modest, positive correlation was observed between the two independent knockdown lines, KD clone 8.1 and KD clone 8.2 (r = 0.39, P < 0.001; **Figure 3A**), consistent with their shared experimental methodology. Notably, the correlation between the KO model and KD clone 8.2 was the strongest of the three comparisons (r = 0.54, P < 0.001; **Figure 3B**), suggesting that despite differences in perturbation strategy, this knockdown clone most closely recapitulates the transcriptional consequences of complete CHD8 loss. Although the fitted regression slope in **Figure 3B** appears visually shallow relative to the line of identity (y = x), this reflects the compressed dynamic range of log_2_ fold-changes in the partial knockdown (KD8.2) relative to the complete knockout, rather than a weak underlying association; the Pearson correlation coefficient already accounts for this scale difference and reflects genuine, statistically robust concordance between the two datasets. While the magnitude of these correlations reflects the expected noise of cross-platform/cross-clone comparisons, the consistent positive directionality across all pairwise comparisons support the interpretation that our transcriptomic signatures capture genuine, if variably penetrant, biological consequences of CHD8 disruption during a short period of time (CHD8 KD) or stable and adapted CHD8 KO cell lines.

**Figure 3.**
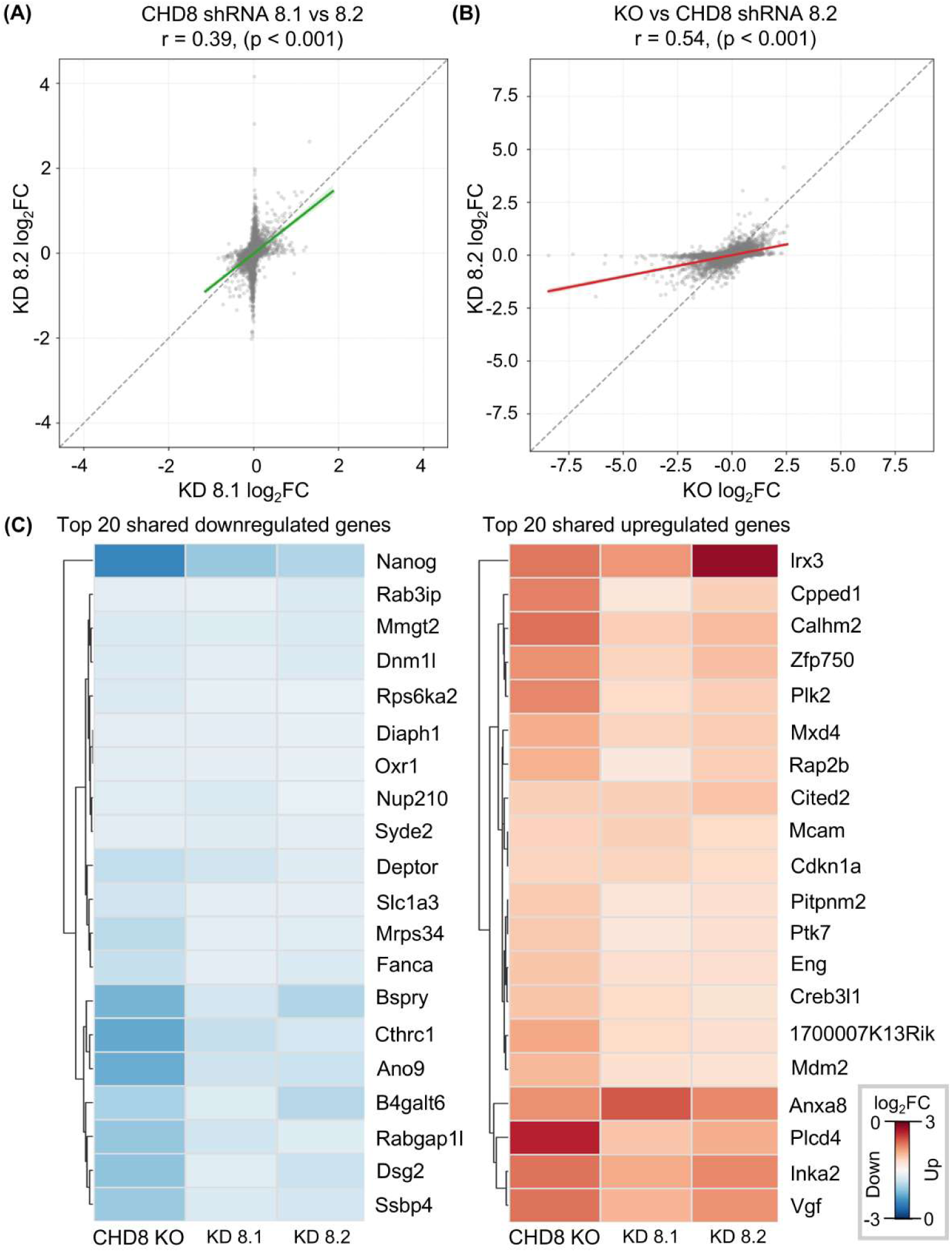
Concordance of gene expression changes across CHD8 perturbation models. **(A, B)** Scatter plots illustrating the correlation of log_2_ fold changes (LFC) between pairs of CHD8 perturbation models. Each point represents an individual expressed gene. Solid colored lines indicate the linear regression fit, with the shaded area representing the 95% confidence interval. The dashed grey diagonal line represents the line of perfect identity (y = x). **(A)** Correlation between Clone 8.1 (KD8.1_LFC) and Clone 8.2 (KD8.2_LFC). A significant positive correlation is observed (r = 0.39, p < 0.001). **(B)** Correlation between the CHD8 knockout model (KO_LFC) and Clone 8.2 (KD8.2_LFC). A significant positive correlation is observed (r = 0.54, p < 0.001). **(C)** Hierarchically clustered heatmaps of the top 20 shared downregulated genes (left) and top 20 shared upregulated genes (right) across all three CHD8 perturbation conditions. Genes shown are significantly deregulated (log_2_FC < 0 or log_2_FC > 0, respectively; DESeq2 adjusted P < 0.05) in CHD8 KO, KD 8.1, and KD 8.2 simultaneously, ranked by mean absolute fold change across conditions. Rows are hierarchically clustered by expression similarity; columns are held in fixed order (KO → KD 8.1 → KD 8.2). Color scale: left panel, log_2_ fold change from −3 (deep blue, strongest downregulation) to 0 (white); right panel, 0 (white) to +3 (deep red, strongest upregulation).

To identify the core set of genes most robustly and consistently regulated by CHD8, we next examined genes significantly dysregulated in the same direction across all three perturbation conditions simultaneously (DESeq2 adjusted P < 0.05, log_2_FC < 0 or > 0 in KO, KD 8.1, and KD 8.2 for down- and up-regulated sets, respectively). Ranking these shared genes by mean absolute log_2_FC across the three conditions, we visualized the top 20 of each set by hierarchical clustering (**Figure 3C**); fold-change magnitude was consistently greatest in the KO, followed by KD 8.2 and KD 8.1, consistent with the dose-dependent relationship described above. Gene ontology enrichment analysis (Enrichr, GO Biological Process, background restricted to genes tested in all three comparisons; **Supplementary Table: S5**) showed that the shared upregulated genes were significantly enriched for DNA damage response and p53-mediated G1 cell cycle arrest signalling (adjusted P = 5.3 × 10⁻⁶), including CDKN1A and MDM2, canonical p53 pathway effectors consistent with previous reports that CHD8 directly restricts P53 stability, trans-activation, and accessibility at P53 target genes in other cell types [76,77]. No significant enrichment was observed among the shared downregulated genes, indicating this set does not converge on a single annotated pathway despite its consistent direction of change across conditions. Hierarchical clustering showed that fold-change magnitude was consistently greatest in the KO, followed by KD 8.2 and KD 8.1, consistent with the dose-dependent relationship described above. Directional analysis of differentially expressed genes revealed a downregulation bias in the KO (1,618 down vs. 1,134 up) and KD 8.2 (233 down vs. 188 up) conditions, consistent with previous reports that CHD8 functions predominantly as a transcriptional activator [78,79]. This pattern was reversed in KD8.1, which showed more upregulated than downregulated genes (43 up vs. 15 down), likely reflecting the small total number of DEGs (n = 58) in this milder knockdown, where a modest number of secondary or compensatory changes can disproportionately affect the up/down balance. Intersection of the 48 CHD8-interacting proteins identified by mass spectrometry in our previously published work [54] with genes significantly downregulated in CHD8-KO NPCs (n = 1,618) identified 18 overlapping genes; significantly more than expected by chance (hypergeometric test, P = 3.1 × 10⁻^5^; expected ≈ 6.6; **Supplementary Figure 3**), dominated by genes encoding ribosomal proteins (14 of 18), alongside WDR5, USP7, SERBP1 and PDHB. This indicates preferential transcriptional downregulation of genes encoding CHD8’s physical interactome upon CHD8 loss, particularly genes encoding ribosomal proteins. Examination of canonical pluripotency and cell-cycle gene panels showed that pluripotency-associated genes (including NANOG, DPPA3, DPPA5A, and ESRRB) were predominantly downregulated rather than activated, while the p53-pathway effectors CDKN1A and MDM2 [80] were consistently upregulated across all three conditions, alongside reduced expression of the proliferation marker Mki67 in KD 8.1 and KD 8.2. Together with the DNA damage response/p53 enrichment noted above, this pattern is more consistent with CHD8-dependent activation of a p53-mediated cell-cycle arrest program than with a block in exit from pluripotency. Because our integrated multi-omics profiling was restricted to a descriptive snapshot at Day 3 of neuronal differentiation, we cannot definitively exclude a kinetic delay in differentiation trajectories in favour of an absolute block. Future time-resolved longitudinal profiling will be necessary to fully map whether the persistent retention of pluripotency markers in CHD8-KO cells represents a permanent fate disruption or a delayed differentiation kinetic.

**Supplementary Figure 3.**
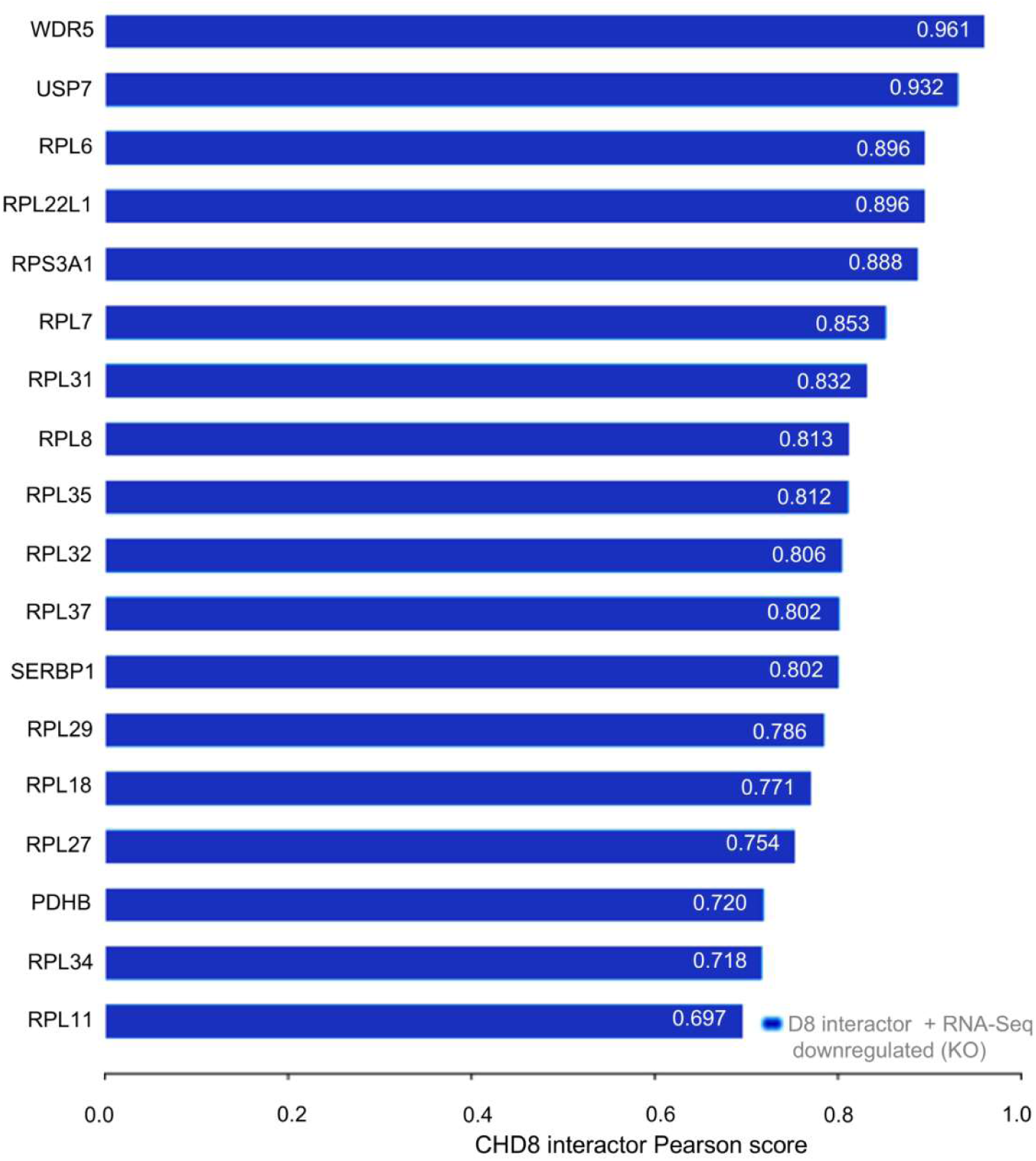
CHD8 interactome–transcriptome overlap in NPCs. Intersection of CHD8-interacting proteins identified by mass spectrometry (Cerase et al., 2021; n = 48, excluding CHD8 itself as bait) with genes significantly downregulated in CHD8-KO NPCs (DESeq2 adjusted *P* < 0.05, |log_2_FC| > 0.5; n = 1,618) identified 18 overlapping genes significantly more than expected by chance (hypergeometric test, P = 5.4 × 10⁻^6^; expected ≈ 5.7; background universe N = 13,590). Gene symbols were standardized prior to matching.The overlap set was dominated by genes encoding ribosomal proteins (14 of 18: RPL6, RPL7, RPL8, RPL11, RPL18, RPL22L1, RPL27, RPL29, RPL31, RPL32, RPL34, RPL35, RPL37, RPS3A1), alongside WDR5, USP7, SERBP1, and PDHB. The bar plot shows all 18 overlapping genes, ranked by mass spectrometry evidence score.

### CHD8 directly regulates neurodevelopmental gene programmes at NPC-specific binding sites and its loss destabilizes a core cohort of highly penetrant Tier 1 ASD risk genes in female progenitors

To identify direct transcriptional targets, we focused on genes directly bound by CHD8 within ± 2 kb of the TSS that were also significantly dysregulated upon CHD8 loss. We focused on a high-confidence set of 254 genes that were both directly bound by CHD8 at NPC-specific peaks [within ± 2 kb of the TSS; derived from 28,690 NPC-specific peaks corresponding to 3,754 unique promoter-proximal target genes, note that this set differs from the ∼2,000 CHD8+H3K4me3 co-occupied genes described above, as it applies a stage-specificity filter (NPC-specific peaks only) rather than a chromatin-state filter (H3K4me3 co-occupancy)] and significantly dysregulated upon CHD8 loss in KO NPCs (DESeq2; adjusted P < 0.05, |log_2_FC| > 0.5; baseMean > 10) of which 109 were upregulated and 145 downregulated. Gene Ontology (GO) Biological Process enrichment analysis of this functionally constrained target set was performed using Enrichr (GO_Biological_Process_2023; organism = mouse), with all expressed genes passing the minimum expression threshold (baseMean > 10) and for which DESeq2 returned an adjusted p-value after independent filtering used as the reference background (n = 13,590). As FDR correction was overly conservative at this list size (n = 254), nominal P < 0.05 was applied, yielding 73 enriched terms of which the top 10 of which are listed in **Supplementary Table: S6**. Although no term survived FDR correction, the top nominally enriched categories converged on two recurring signalling themes rather than a single dominant pathway: G-protein-coupled receptor/adenylate cyclase signalling (ADCY2, ADGRG1, ADGRL1, CALCR, GLP1R, GNA15; P = 8–9 × 10⁻⁴) and TGF-β/SMAD signalling (TGFB1, SMAD6, BMP8B, PMEPA1), with TGFB1 recurring across both the SMAD-phosphorylation and cardiac muscle morphogenesis terms. A smaller cluster of genome-defense genes (SIRT6, TDRD12, TUT7) was enriched for retrotransposon silencing, the single most significant term overall (P = 7.0 × 10⁻⁴). Given the modest list size and lack of FDR-significant terms, these associations should be considered hypothesis-generating rather than definitive evidence of a coordinated CHD8-dependent transcriptional program. To further summarise these results, we grouped enriched terms into four major biological modules identified by keyword matching within GO results: Synaptic function, Chromatin organisation, Axon development, and Transcriptional regulation (**Figure 4A**). Importantly, this focused analysis of directly regulated NPC-specific CHD8 supports a model in which CHD8 regulates neuronal transcriptional programmes at direct target loci during differentiation.

**Figure 4.**
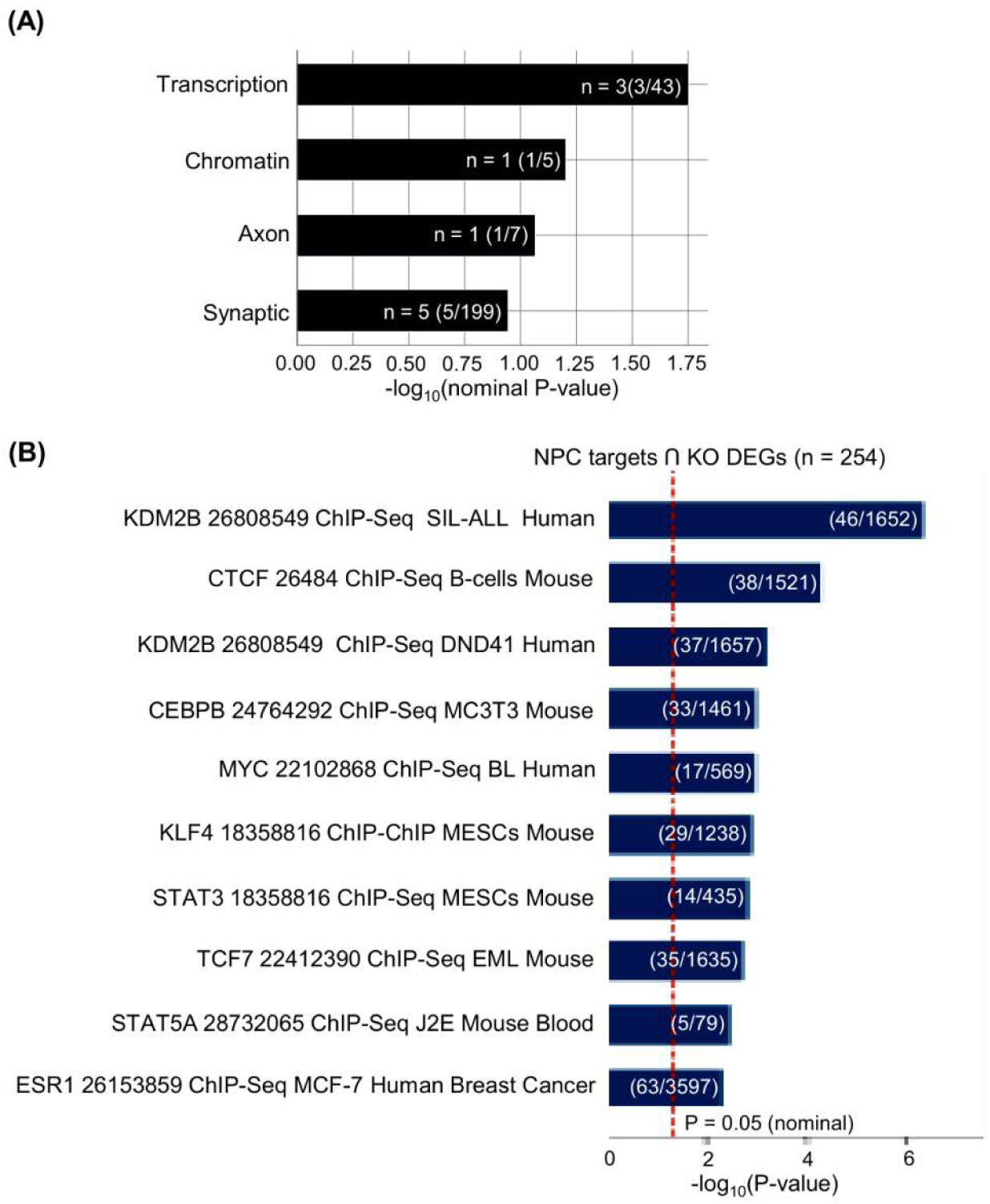
Functional characterisation of NPC-specific CHD8 target genes. **(A)** Bar plot summarising major functional modules associated with NPC-specific CHD8 targets. Representative genes from key categories (Synaptic, Chromatin, Transcription, and Axon-related processes)enriched among the 254-gene high-confidence CHD8 target set, ranked by the significance of each module’s representative GO Biological Process term. Bars are annotated with the number of genes and the term’s overlap fraction. **(B)** Transcription factor co-occupancy analysis for the same 254 gene high confidence CHD8 target set. Bar plot showing the top enriched transcription factor binding signatures, suggesting co-regulatory networks involving CHD8 and factors such as KDM2B, CTCF, and MYC.

To place these targets in a broader regulatory context, we performed transcription factor co-occupancy analysis using ChEA_2022 on the same 254-gene input set, identifying 53 significant TF associations (nominal P < 0.05), of which the top 10 are shown (**Figure 4B**, **Supplementary Table: S7**). This revealed significant enrichment for binding signatures of core architectural and regulatory factors, including KDM2B, CTCF, CEBPB, MYC, KLF4, STAT3, TCF7, and ESR1. We note its established cooperative role of Chd8 with CTCF in coordinating structural promoter-enhancer interactions [17,81]. Together, these enrichments strongly suggest that CHD8 operates within a highly coordinated chromatin and transcriptional network rather than acting in isolation, consistent with roles in chromatin organisation, promoter architecture, and transcriptional control in neural progenitor cells.

To assess the relationship between our female multi-omics framework and clinical ASD genetic vulnerability, we mapped the 2,752 significantly altered DEGs to the curated SFARI Autism Risk Gene dataset. Out of 905 SFARI-annotated genes expressed in our baseline progenitor population (N = 13,590), 185 genes overlapped with our dysregulated gene set. Of the 185 overlapping genes, 165 were classified as SFARI Tier 1 (n = 41, high-confidence), Tier 2 (n = 88, strong candidate), or Tier 3 (n = 36, suggestive evidence); the remaining 20 were annotated as SFARI syndromic, including genes linked to known monogenic disorders such as Duchenne muscular dystrophy (DMD) and Lowe syndrome (OCRL). Across the full 185-gene set **(Supplementary Table: S8)**, 107 genes were downregulated and 78 upregulated. This cohort includes established neurodevelopmental hubs such as EBF3, PTCHD1, AUTS2, and SCN1A, alongside regulators including MECP2, FMR1, and CDKL5, all of which were downregulated **(Figure 5)**.

**Figure 5.**
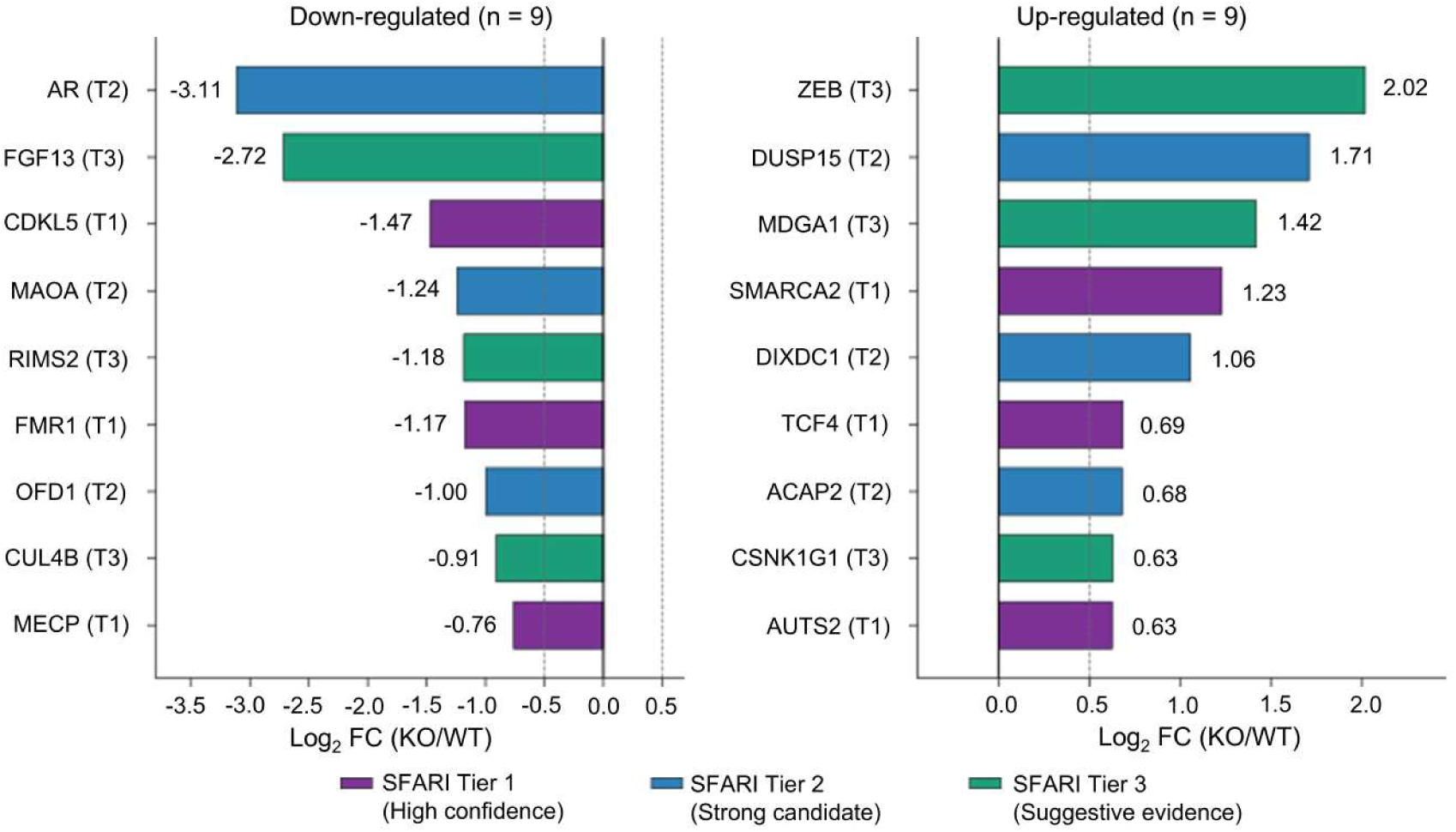
Transcriptional dysregulation of representative, well-characterized autism spectrum disorder (ASD) risk genes spanning SFARI confidence Tiers 1–3 in CHD8-KO NPCs. Two-panel horizontal bar plot showing log_2_ fold change (FC, KO/WT) for a curated, representative set of ASD risk genes spanning SFARI Tier 1 (high confidence), Tier 2 (strong candidate), and Tier 3 (suggestive evidence), following CRISPR-mediated CHD8 homozygous knockout. Of 165 significantly dysregulated Tier 1-3 genes identified in this dataset (92 downregulated, 73 upregulated; see Results), 9 representative genes per direction are shown here, prioritizing well-established ASD genes; the complete gene list is provided in Supplementary Table: S8. **(A)** Downregulated genes (n = 9 shown): Tier 1 (CDKL5, FMR1, MECP2), Tier 2 (AR, MAOA, OFD1), and Tier 3 (FGF13, RIMS2, CUL4B). **(B)** Upregulated genes (n = 9 shown): Tier 1 (SMARCA2, TCF4, AUTS2), Tier 2 (DUSP15, DIXDC1, ACAP2), and Tier 3 (ZEB2, MDGA1, CSNK1G1). Within each panel, genes are ordered by absolute magnitude of log_2_FC relative to wild-type. Bar fill color denotes SFARI confidence Tier (purple = Tier 1; blue = Tier 2; green = Tier 3), as indicated in the shared legend below the panels. Numerical labels adjacent to each bar report exact log_2_FC values calculated via DESeq2. Dashed vertical lines mark the ± 0.5 log_2_FC selection threshold. All displayed genes met row-filtering criteria (baseMean > 10) and statistical significance (FDR < 0.05).Note that the overall overlap between CHD8-KO DEGs and SFARI-annotated genes did not exceed chance expectation genome-wide (see Results); genes shown here are highlighted for their established relevance to ASD, not as evidence of statistical enrichment.

### Chromatin accessibility changes reveal CHD8’s role in establishing neuronal epigenetic landscapes

To determine whether CHD8 loss is accompanied by global changes in chromatin accessibility, we reanalyzed ATAC-seq on WT and CHD8-KO NPCs (originally generated in prior study [54]) using DiffBind and DESeq2. A consensus peak set was constructed from four samples (two WT replicates and one replicate from each of two independent CHD8-KO clones), requiring peak support in at least two of the four samples irrespective of genotype (minOverlap = 2, no condition-specific masking), so that both shared and genotype-specific accessible regions were retained rather than restricting analysis to peaks common to WT and KO alone. Prior to this, principal component analysis identified one KO replicate (KO1_rep1) as a clear outlier inconsistent with its assigned clone and condition; this sample was excluded from all downstream analyses, and read counting and differential testing were performed across the five remaining non-outlier replicates (two WT, one KO clone-1, two KO clone-2) against the four-sample consensus set, yielding 74,486 coordinate-matched regions.

Analysis of log_2_ fold change values across all 74,486 regions revealed that CHD8 loss results in a negligible net shift in global chromatin accessibility (median LFC = +0.014), with 51.6% of regions (38,454) showing increased accessibility and 48.4% (36,032) showing decreased accessibility in CHD8-KO relative to WT NPCs (**Figure 6A**, **Supplementary Table: S9**), rather than a global reduction consistent with CHD8 playing a role in both maintaining and restraining chromatin accessibility at distinct loci [82].

**Figure 6.**
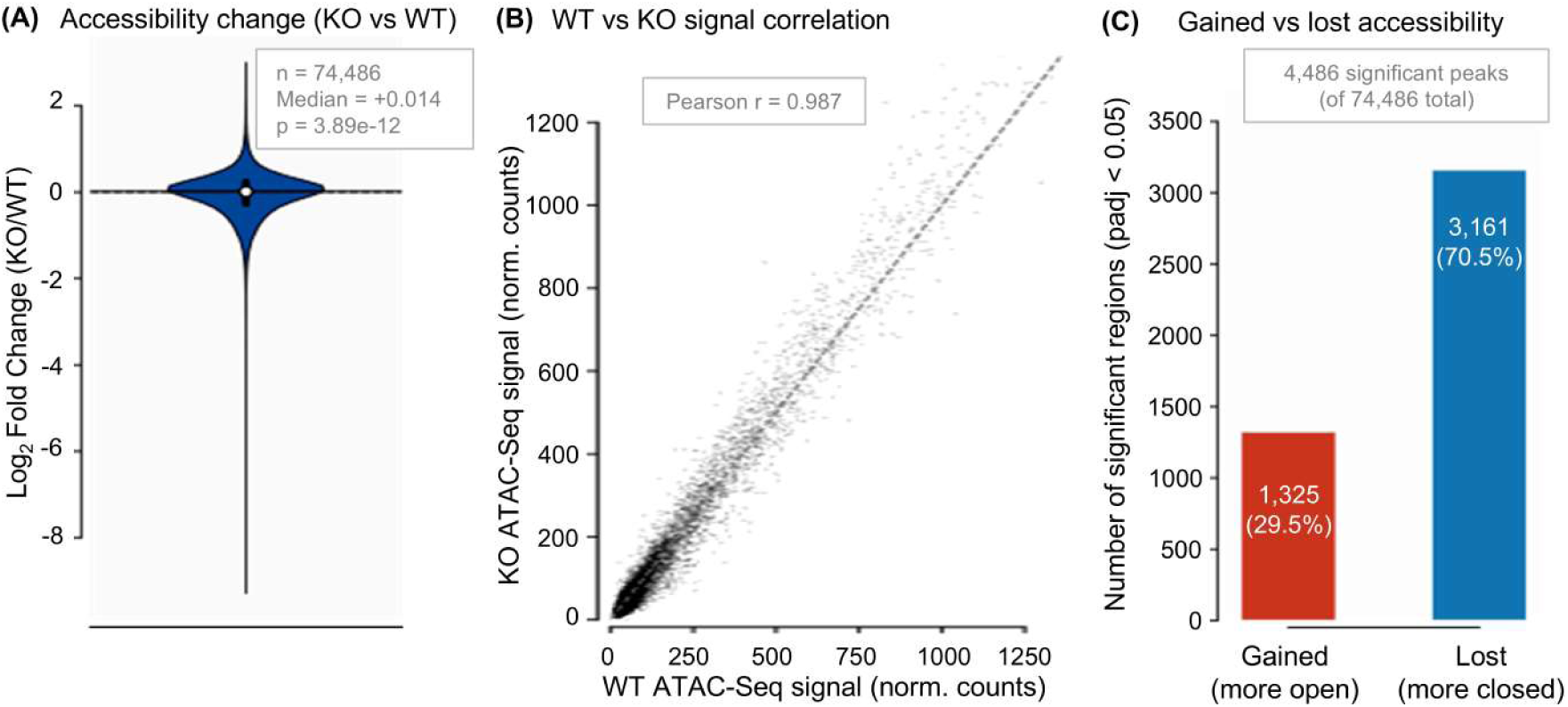
Chromatin accessibility changes following CHD8 loss in neural progenitor cells. **(A)** Global distribution of chromatin accessibility changes in CHD8 knockout (KO) versus wild-type (WT) cells. Violin plot showing the distribution of log_2_ fold change (LFC) in ATAC-seq signal across 74,486 consensus regions, identified using DiffBind (minOverlap = 2 across four samples: two WT replicates and one replicate from each of two independent CHD8-KO clones, irrespective of genotype) and tested for differential accessibility by DESeq2 across five non-outlier replicates (one KO replicate excluded as a confirmed PCA outlier). Overall, 51.6% of regions exhibit increased accessibility in KO, while 48.4% show decreased accessibility. The median LFC (+ 0.014) indicates a negligible net shift in global chromatin accessibility upon CHD8 loss. The dashed red line denotes zero change (parity). **(B)** Correlation of chromatin accessibility between WT and CHD8-KO cells at matched genomic regions. Scatter plot of DESeq2-normalizedATAC-seq signal intensities (WT vs KO) across all 74,486 regions. The dashed red diagonal represents perfect concordance (y = x). A Pearson correlation coefficient (r = 0.987) indicates strong global concordance between conditions, consistent with accessibility changes being confined to a comparatively small subset of loci rather than reflecting widespread remodelling. **(C)** Summary of gained and lost chromatin accessibility among statistically significant regions in CHD8-KO cells. Bar plot showing the number and proportion of significant regions (padj < 0.05, |LFC| > 0.5; n = 4,486) with increased (gained; 1,325; 29.5%) or decreased (lost; 3,161; 70.5%) accessibility relative to WT. Regions were classified based on the direction of log_2_ fold change among those passing significance and effect-size thresholds.

Correlation analysis of DESeq2-normalized ATAC-seq signal intensities between WT and CHD8-KO NPCs revealed markedly strong global concordance between conditions (Pearson r = 0.987), with the vast majority of regions showing highly similar signal between genotypes (**Figure 6B**, **Supplementary Table: S9**). This indicates that CHD8 loss does not cause wholesale chromatin compaction but instead produces locus-specific accessibility changes concentrated at specific regulatory elements.

Consistent with the balanced global distribution observed in panel A, restricting to regions reaching statistical significance (padj < 0.05 and |LFC| > 0.5, n = 4,486) revealed a clear predominance of accessibility loss in CHD8-KO NPCs, with 3,161 regions (70.5%) losing and 1,325 regions (29.5%) gaining accessibility relative to WT (**Figure 6C**, **Supplementary Table: S9**). This contrasts with the near-even split seen across all regions irrespective of significance, indicating that although CHD8 loss does not shift the bulk chromatin landscape in either direction, the changes that are statistically robust are predominantly losses of accessibility — consistent with CHD8 acting primarily to maintain accessibility at a defined subset of loci.

To place these chromatin accessibility changes in a broader regulatory context, we performed multi-omics integration of RNA-seq, CHD8 ChIP-seq, and ATAC-seq datasets. Differentially expressed genes (DEGs; padj < 0.05, |LFC| > 0.5) numbered 2,752 (1,134 up, 1,618 down); CHD8 ChIP-seq identified 3,754 promoter-proximal target genes; and significant ATAC-seq regions (padj < 0.05, |LFC| > 0.5; n = 4,486) mapped to 473 genes within 2 kb of a TSS. Three-way overlap analysis identified 8 genes common to all three datasets — simultaneously CHD8-bound, transcriptionally dysregulated, and associated with a significant ATAC-seq accessibility change in KO cells — representing the highest-confidence set of direct CHD8 targets linking chromatin occupancy, accessibility, and transcriptional regulation (**Supplementary Figure 1**). The remaining intersections were partitioned as RNA-only (2,476), ChIP-only (3,346), ATAC-only (289), RNA ∩ ChIP (246), RNA ∩ ATAC (22), and ChIP ∩ ATAC (154), illustrating the relative contributions and degree of overlap between the three regulatory layers. Together these findings indicate that CHD8 loss produces predominantly locus-specific accessibility changes — balanced in aggregate but skewed toward loss among high-confidence regions — and that this locus-specific pattern converges with transcriptional dysregulation and CHD8 binding at a small, stringently defined set of 8 neurodevelopmental target genes, though the limited size of this high-confidence set (SLC17A9, DSCC1, TFAP2C, BMP8B, ABHD4, INPP5E, P2RX3, CREB3L1) precluded formal statistical enrichment testing. Gene function analysis identified two genes directly involved in purinergic (ATP) signalling (SLC17A9, P2RX3), consistent with a potential role for CHD8 in neuronal signalling pathways, alongside developmental signalling regulators (BMP8B, INPP5E) that echo the TGF-β/SMAD and GPCR-associated themes identified in the broader 254-gene target set **(Figure 4A)**. To identify transcription factors potentially co-operating with CHD8 at Broad-set loci, we performed TF co-binding enrichment analysis on the 254 Broad-set genes, querying the ENCODE TF ChIP-seq, ChEA 2022, ENCODE + ChEA Consensus, TRANSFAC/JASPAR, and TF Perturbations databases via Enrichr. TF perturbation-based analysis identified three signatures significantly enriched among Broad-set genes (all FDR = 0.024): knockdown of KDM2B (15/327 genes), knockout of NFE2L2 (17/419 genes), and overexpression of ARID3A (15/346 genes). Independently, ChEA 2022 ChIP-seq enrichment identified KDM2B binding (46/1,652 genes, FDR = 2.4 × 10⁻⁴) and CTCF binding (38/1,521 genes, FDR = 0.017) as significantly enriched among Broad-set gene promoters. No terms reached FDR < 0.05 in the ENCODE TF ChIP-seq 2015, ENCODE + ChEA Consensus, or TRANSFAC/JASPAR databases. The recurrence of KDM2B across two independent database types (perturbation-response signatures and direct ChIP-seq binding), together with ARID3A and CTCF enrichment, is consistent with CHD8 cooperating with Polycomb-associated and architectural chromatin regulators at target promoters. Collectively, these results suggest that CHD8 may cooperate with a small number of chromatin regulators and sequence-specific transcription factors — including KDM2B, ARID3A, NFE2L2, and CTCF — to modulate transcriptional programs at Broad-set target genes (**Supplementary Figure 4, Supplementary Table: S10-14**).

**Supplementary Figure 4.**
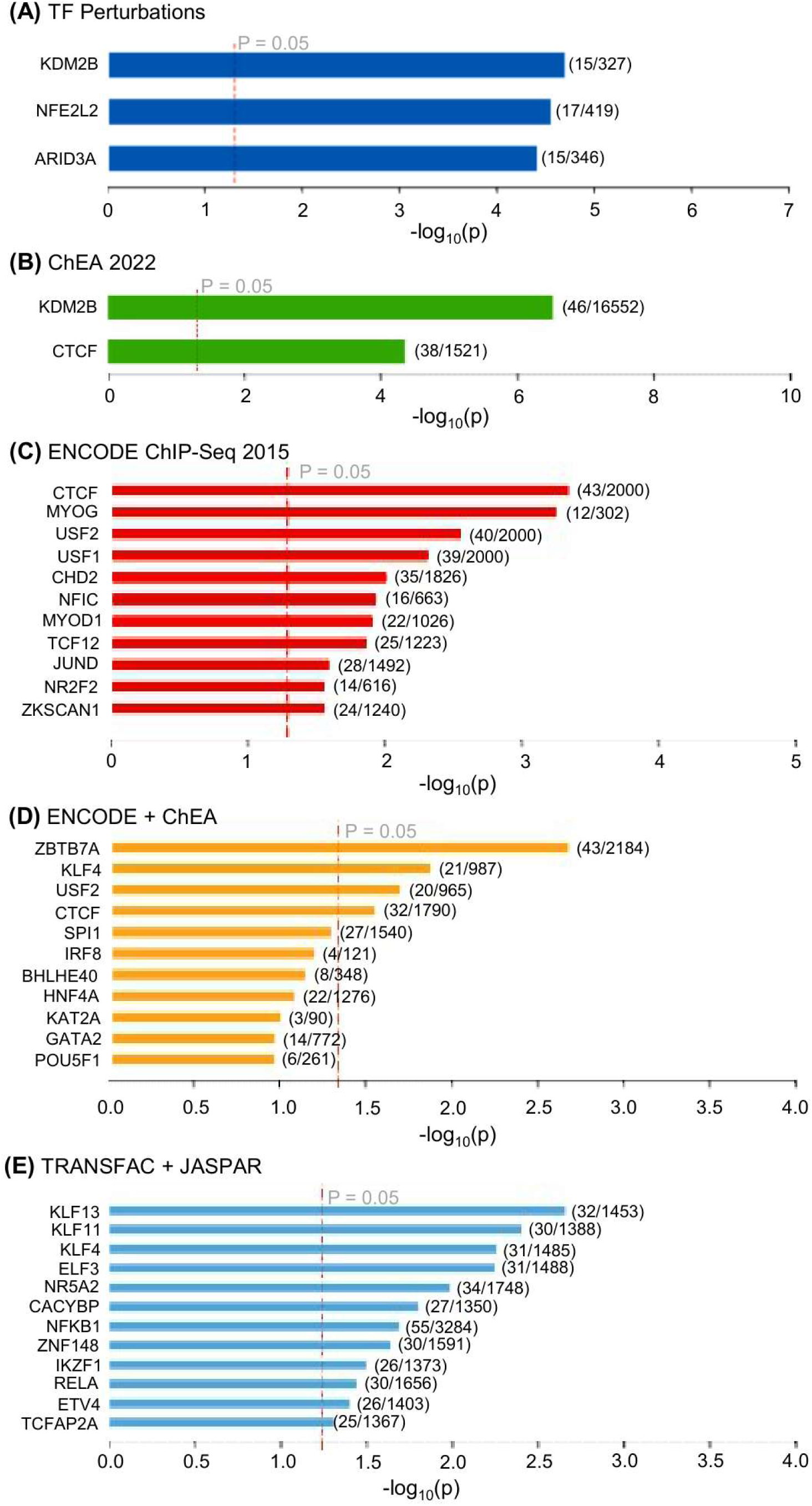
Transcription factor enrichment analysis of CHD8-associated target genes. (Broad set, n = 254). TF co-binding enrichment analysis was performed on the 254 genes comprising the Broad set (CHD8-bound + significantly differentially expressed, independent of ATAC-seq significance; see Methods). Enrichment was assessed across multiple curated databases via Enrichr: **(A)** TF Perturbations, **(B)** ChEA 2022, **(C)** ENCODE TF ChIP-seq 2015, **(D)** ENCODE + ChEA Consensus, and **(E)** TRANSFAC/JASPAR. Bar plots show the top enriched terms ranked by −log10(p-value); the red dashed line indicates the nominal significance threshold (p = 0.05). TF Perturbations analysis identified three signatures reaching FDR < 0.05: KDM2B knockdown, NFE2L2 knockout, and ARID3A overexpression (all FDR = 0.024). ChEA 2022 independently identified KDM2B and CTCF ChIP-seq binding as significantly enriched (FDR = 2.4 × 10⁻⁴ and 0.017, respectively). No terms reached FDR < 0.05 in the remaining three databases.

### Chromodomain and helicase domain make distinct contributions to CHD8-mediated transcriptional rescue

To dissect the functional contributions of CHD8’s chromodomains and helicase domain to transcriptional rescue, we generated CHD8-KO NPCs re-expressing full-length CHD8 (FL), chromodomain-deleted CHD8 (ΔChromo), or helicase-deleted CHD8 (ΔHelicase). The domain architecture and specific deletions for each construct are illustrated in **Supplementary Figure 2**. We confirmed by Western blot and next-generation sequencing both the loss of CHD8 protein in the C4.1 KO line and successful re-expression of FL, ΔChromo, and ΔHelicase variants in the KO background at comparable levels (**Supplementary Figure 2**). Rescue efficiency was then assessed across NPC-specific direct CHD8 target genes with |log_2_FC| > 0.5 in CHD8-KO cells. Rescue fraction (RF) was calculated for each gene as RF = (LFC_rescue − LFC_KO) / (0 − LFC_KO), where RF = 1 indicates complete restoration to wild-type expression levels and RF = 0 indicates no rescue (**Figure 7A**, **Supplementary Table: S15**). Full-length CHD8 achieved the highest median rescue fraction among the three constructs, indicating that the intact protein is required to achieve the strongest transcriptional rescue; however, rescue by FL was partial (∼ 70%) rather than complete (median RF < 1), as expected. Notably, this partial rescue is not attributable to inadequate re-expression, as DESeq2-normalised CHD8 expression in the FL rescue line did not differ significantly from CTRL **(Supplementary Figure 2C)**, indicating that FL protein was restored to wild-type levels. This observation is consistent with CHD8 not acting as a pioneer factor at many loci, including a subset of X-linked genes that remained un-rescued, and this partial rescue does not on its own exclude a contribution from off-target effects of the knockout or re-expression system. Deletion of either domain substantially impaired rescue capacity in distinct ways. ΔChromo exhibited no recovery (∼ 1%), indicating that chromodomain integrity is essentially indispensable for CHD8-mediated transcriptional rescue. ΔHelicase showed partial rescue (∼ 41%), demonstrating that helicase catalytic activity makes a substantial and non-redundant contribution, though a proportion of target gene expression can be partially restored in its absence. Statistical comparison confirmed significant differences in rescue fractions across constructs by Friedman test, with all pairwise Wilcoxon signed-rank comparisons reaching significance after FDR correction (**Figure 7A**).

**Figure 7.**
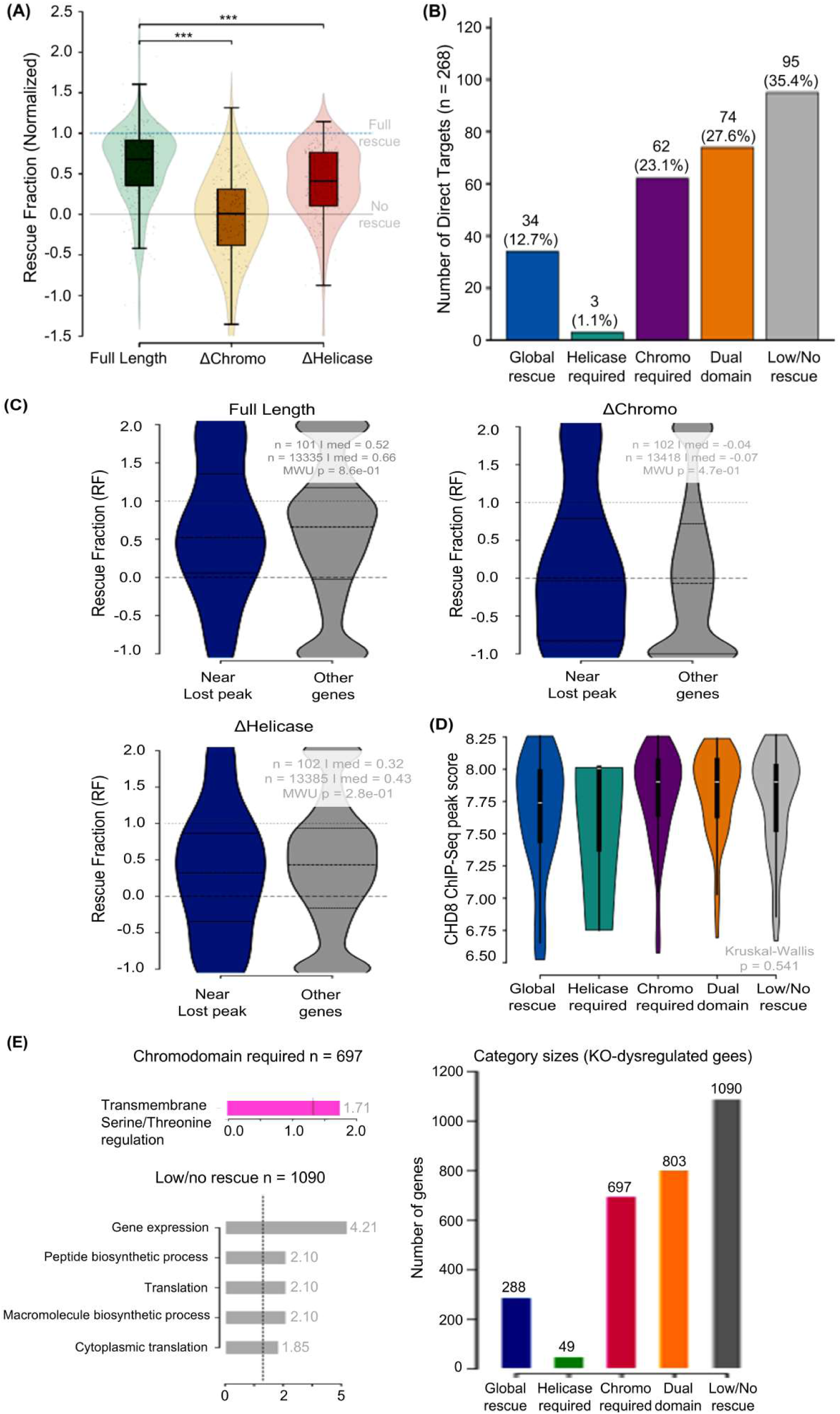
Domain-dependent requirements of CHD8 for transcriptional rescue of NPC target genes. **(A)** Domain-specific rescue efficiency across direct CHD8 target genes in NPCs. Violin–box– strip plots depict the distribution of rescue fraction (RF) values for each re-expression construct, calculated for genes with |log_2_FC| > 0.5 in CHD8-KO cells. RF is defined as (LFC_rescue – LFC_KO)/(0 – LFC_KO), where RF = 1 (blue dashed line) indicates complete restoration to wild-type levels and RF = 0 (black line) indicates no rescue. Full-length CHD8 shows the highest median rescue, whereas ΔChromo exhibits minimal recovery and ΔHelicase shows partial rescue. Violin shapes represent density, box plots indicate median and interquartile range, and points show individual genes. Statistical significance was assessed using a Friedman test followed by pairwise Wilcoxon signed-rank tests with FDR correction. **(B)** Classification of CHD8 target genes based on domain-specific rescue patterns. Genes were grouped into five mutually exclusive categories using an RF threshold of ≥ 0.5: globally rescued, helicase-dependent, chromodomain-dependent, dual-domain-dependent, and low/no rescue. The uneven distribution across categories (χ^2^ test) indicates that distinct subsets of CHD8 targets require different structural domains for effective transcriptional recovery. **(C)** Relationship between chromatin accessibility loss and rescue efficiency. Violin plots compare RF values for genes proximal to significant difference in RF between near-lost-peak and other genes (Mann–Whitney U test: FL p = 0.86, ΔChromo p = 0.47, ΔHelicase p = 0.28), and the direction of the modest observed differences was not consistent across constructs. Overall, proximity to significant chromatin accessibility loss does not strongly or consistently predict domain-specific rescue outcome in the current dataset. **(D)** CHD8 binding intensity and occupancy across domain-requirement categories. Violin plots showing the distribution of CHD8 ChIP-seq peak scores (log₁₀) at gene-proximal loci (within 2kb of TSS) for each category, calculated across all KO-dysregulated genes (n = 2,927; |log_2_FC| > 0.5, baseMean > 10). Bar plot showing the percentage of genes with a CHD8 peak within 2 kb of the TSS per category. All five categories show equivalent CHD8 occupancy (100%) and comparable binding intensity (Kruskal–Wallis test), indicating that differential CHD8 recruitment does not explain domain-requirement category membership. **(E)** Gene Ontology Biological Process enrichment analysis for domain-requirement categories. Bar plots show significantly enriched GO terms (FDR < 0.05) identified by Fisher’s exact test with Benjamini– Hochberg correction, using all expressed KO-dysregulated genes as background. Left: chromodomain-dependent genes are enriched for regulation of transmembrane receptor serine/threonine kinase signalling (GO:0090092; FDR = 0.02), consistent with a requirement for precise chromodomain-mediated targeting at developmentally regulated signal-responsive loci. Right: low/no rescue genes are enriched for core biosynthetic and translational processes including cytoplasmic translation, peptide biosynthesis, and gene expression (FDR < 0.05), consistent with constitutively expressed housekeeping genes refractory to CHD8 re-expression. Categories without FDR-significant enrichment (globally rescued, helicase-dependent, dual-domain-dependent) likely reflect functional heterogeneity within these groups. Gene counts per category: Global Rescue n = 288, Helicase required n = 49, Chromodomain required n = 697, Dual Domain required n = 803, Low/No Rescue n = 1,090.

Categorical classification of CHD8 direct NPC target genes by construct-specific rescue (RF ≥ 0.5 threshold) revealed a non-uniform distribution across five mutually exclusive rescue categories: global rescue (rescued by all three constructs), helicase motor required (rescued by FL and ΔChromo but not ΔHelicase), chromodomain required (rescued by FL and ΔHelicase but not ΔChromo), both domains required (rescued by FL only), and low/no rescue (not rescued by any construct) (**Figure 7B**, **Supplementary Table: S16**). The non-uniform distribution across these categories was confirmed by chi-square goodness-of-fit test (χ^2^ = 95.99, p = 7.01 × 10⁻²⁰), establishing that different CHD8 target genes have distinct and separable domain requirements for transcriptional rescue.

To examine whether chromatin accessibility changes at KO-lost ATAC-seq peaks influence domain-specific rescue efficiency, we compared rescue fractions between genes proximal to KO-lost peaks (n = 347 unique genes within 2 kb of a significant KO-lost peak TSS; padj < 0.05, LFC < −0.5) and all other genes across all three constructs (**Figure 7C**, **Supplementary Table: S17**).

While some directional differences were observed among genes proximal to significant KO lost peaks (n = 347) — modestly lower median RF for FL (median 0.52 vs. 0.66) and ΔHelicase (median 0.32 vs. 0.43) and a modestly higher median RF for ΔChromo (−0.04 vs. −0.07) — none of these differences reached statistical significance (FL p = 0.86, ΔChromo p = 0.47, ΔHelicase p = 0.27), and the direction of the effect was not consistent across constructs. This indicates that proximity to significant KO-lost chromatin accessibility regions does not reliably stratify genes into domain-dependent rescue groups. This finding is further reinforced by analysis of CHD8 binding as a non-predictor of rescue: of 1,476 direct CHD8 targets detected in the RNA-seq dataset, 856 (58.0%) were rescued by FL re-expression, comparable to the 57.2% rescue rate observed among indirect DEGs (n = 11,910; Fisher’s exact test: OR = 1.03, p = 0.558; **Supplementary Figure 5A, B**). Consistently, the distribution of RF values was similar between direct CHD8 targets and non-target genes (Mann–Whitney U test: p = 0.106; **Supplementary Figure 5C**), confirming that CHD8 binding alone does not stratify genes by rescue efficiency. Together these findings indicate that transcriptional rescue by CHD8 involves complex multi-factor regulation that extends beyond chromatin accessibility or binding occupancy alone.

**Supplementary Figure 5.**
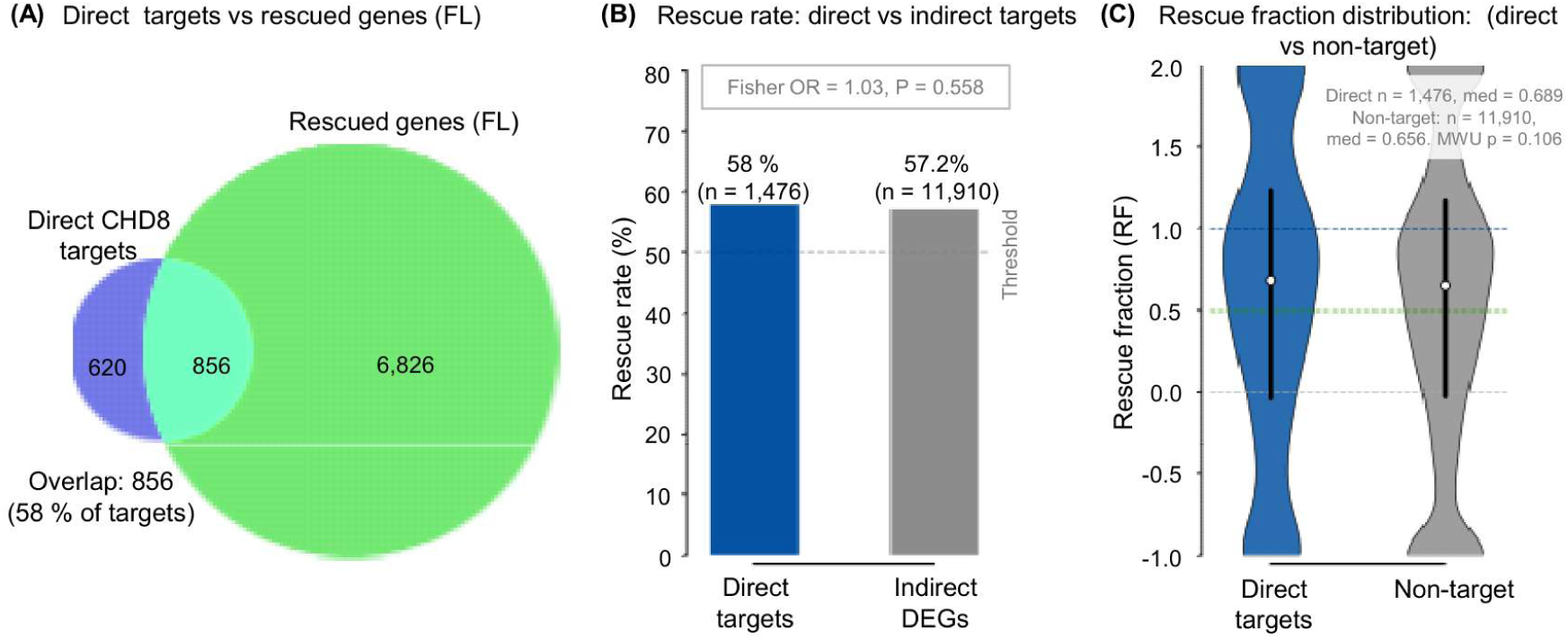
CHD8 binding does not predict full-length rescue efficiency. Assessment of whether direct CHD8 ChIP-seq target genes show preferential rescue upon full-length (FL) CHD8 re-expression in NPCs. Direct targets were defined as genes with a CHD8 peak within 2 kb of the TSS in NPCs (n = 3,754; n = 1,476 detected in RNA-seq after baseMean > 10 filtering). Rescue fraction (RF) was calculated per gene, and genes with RF ≥ 0.5 were classified as rescued. **(A)** Overlap between direct CHD8 targets (n = 1,476) and FL-rescued genes (n = 7,682) shows that 856 targets (58.0%) are rescued. **(B)** Rescue rates are comparable between direct targets (58.0%) and indirect DEGs (57.2%; n = 11,910), with no significant enrichment (Fisher’s exact test: OR = 1.03, p = 0.558). **(C)** Distribution of RF values is similar between direct targets and non-target genes (Mann–Whitney U test: p = 0.106), indicating that CHD8 binding alone does not stratify genes by rescue efficiency.

## Discussion

### CHD8 re-localises from pluripotency to neuronal loci during female neuronal differentiation

Our multi-omics analysis identifies CHD8 as a key regulator of female neuronal differentiation. By binding H3K4me3-marked promoters and remodelling chromatin in an ATP-dependent manner, CHD8 coordinates repression of pluripotency programmes, activation of neurogenic genes, and expression of synaptic gene networks. Comparing CHD8 occupancy between ES cells and NPCs revealed extensive redistribution of the cistrome during differentiation: the majority of ES-cell peaks are vacated, while 28,690 NPC-specific peaks emerge, corresponding to 3,754 genes that gain promoter-proximal CHD8 binding only after neural commitment. This shift from stem-cell regulators toward neuronal genes indicates that CHD8 is dynamically repositioned as cells exit pluripotency and adopt a neural fate.

In female NPCs, only 31.7% of CHD8 binding events occur at promoters, with the majority (66.6%) located at distal regulatory elements (**Figure 1B**). Whether this distribution reflects sex- or species-specific features, or simply methodological differences in how the CHD8 cistrome has been mapped across studies, will require matched male-female comparisons using identical pipelines. Despite this uncertainty, the transcription factor co-occupancy landscape is conserved: CTCF is strongly enriched at CHD8 peaks in our dataset (**Figure 4A**), consistent with earlier reports in male-derived human NPC and mouse ESC/NPC systems identifying a CTCF motif as the most significantly enriched sequence at CHD8 binding sites [17], independently supported by CHD8 ChIP-seq studies in human fetal brain [16] and murine ESC-derived neural progenitors [75], pointing to preserved core regulatory partnerships across systems and species. At the transcriptional level, CHD8 loss produced a marked directional asymmetry, with downregulated genes outnumbering upregulated ones in the knockout line (1,618 down vs. 1,134 up) and in the stronger knockdown line (KD2: 233 down vs. 188 up), whereas the milder knockdown (KD1) showed the opposite pattern (43 up vs. 15 down), likely reflecting its smaller number of total DEGs and reduced knockdown efficiency at this locus (**Figure 2A**). A limitation of this study is that our CHD8-KO experiments rely on a single CRISPR-derived clone (C4), so clone-specific off-target or selection effects cannot be formally excluded. However, the significant positive correlation between the KO transcriptome and the independently-derived KD8.2 knockdown line (r = 0.54, p < 0.001), together with the substantial transcriptional rescue achieved by full-length CHD8 re-expression in the same KO background (∼70% of dysregulated targets), both indicate that the observed KO phenotype reflects genuine CHD8-dependent regulation rather than a clone-specific artefact. We also note that the parental Fa2L-S4 line carries a Tsix mutation, which biases the choice of which X chromosome is inactivated during XCI but does not otherwise alter neuronal differentiation or CHD8 function. CHD8 binding itself, however, was a poor predictor of transcriptional outcome: bound and unbound genes showed no overall difference in expression change (**Figure 2C**), implying that most CHD8-dependent transcriptional effects might be indirect, in agreement with previous work [16,17]. To ask whether this transcriptionally silent binding still has chromatin consequences, we examined the 3,500 CHD8-bound genes that showed no differential expression after CHD8 loss and asked whether they nonetheless underwent significant accessibility changes. 4.4 % did, a proportion not significantly different from the 3.2 % seen among CHD8-bound genes that were differentially expressed (odds ratio = 0.71, Fisher’s exact p = 0.42). This equivalence suggests that CHD8’s chromatin-remodelling activity at a bound locus is largely uncoupled from whether that locus changes expression, though such accessibility changes are a comparatively rare event overall – occurring at only a small minority of CHD8-bound genes regardless of transcriptional status - with transcriptional consequences, where they occur, likely emerging only where additional regulatory inputs (co-factor availability, enhancer connectivity, or signal-dependent activation) are also engaged. Together, these observations support a model in which CHD8’s accessibility-maintaining activity at bound loci is real but restricted to a defined subset of targets, rather than a pervasive property of binding itself.

### CHD8 targets converge on ASD-risk pathways

Pathway enrichment among genes downregulated in CHD8-KO NPCs highlighted suppression of cytoplasmic translation and ion-transport processes (**Supplementary Table: S2**).Together, these findings indicate that CHD8 loss disrupts both neuronal differentiation programmes and the biosynthetic machinery that supports them, consistent with the high penetrance of CHD8-associated ASD and with dysregulation of convergent pathway components extending to female neuronal cells. CHD8 targets co-occupied by KDM2B, CTCF, and MYC (**Figure 4B**, **Supplementary Table: S7**) point to CHD8 acting within coordinated chromatin-regulatory networks rather than in isolation, consistent with a model in which CHD8 loss disrupts entire gene-expression modules rather than individual genes. Cross-referencing our dataset against the SFARI autism risk gene list showed that CHD8 depletion in female NPCs alters expression of 41 Tier 1 high-confidence ASD risk genes (and 88 and 36 Tier 2/3 genes, respectively). Tier 1 genes were enriched for X-linked (15 of 41), most plausibly reflecting the modest – yet significant - Xist upregulation already detectable at day 3 of differentiation; this imbalance may not persist at later developmental stages, as counter-selection could correct it, and it may capture only an early skew in the XCI initiation programme [54]. Even so, this female-derived dataset begins to address the pronounced clinical sexual dimorphism of ASD, though fully resolving the question will require matched, parallel male-female comparisons (see below). Our female NPCs showed a substantial transcriptional response to CHD8 loss (2,752 DEGs), exceeding by approximately 5-fold the 510 DEGs (FDR < 0.05) reported across pooled developmental stages in heterozygous CHD8+/del5 mouse brain [18,19]. Because these studies differ in platform, developmental stage, tissue type, zygosity (heterozygous vs. homozygous knockout), sequencing depth, and analytical approach (individual-gene DEG calling vs. GSEA-based detection of cumulative pathway-level shifts), this difference cannot be interpreted as a direct quantitative sex comparison.

### Both the chromodomain and helicase domains are required for CHD8 function

Our domain-specific rescue experiments clarify how CHD8’s two functional modules contribute to its activity. Full-length CHD8 achieved the highest median rescue fraction across direct NPC target genes, whereas the chromodomain-deletion mutant (ΔChromo) showed minimal recovery, indicating that an intact chromodomain is essentially indispensable for CHD8-mediated transcriptional rescue (**Figure 7A**). This is consistent with CHD8 localisation depending primarily on chromodomain-H3K4me3 interactions, in line with structural work showing high-affinity binding between CHD8 chromodomains and H3K4me3-modified histone tails [83], and with the predominant association we observe between CHD8 occupancy and H3K4me3-marked active promoters in NPCs (**Figure 1B, D**).

The helicase domain also proved important for transcriptional rescue, though its requirement was less abundant than that of the chromodomains: the helicase-deletion mutant (ΔHelicase) achieved only partial rescue, showing that recruitment to chromatin alone is not sufficient - CHD8 must also remodel chromatin structure to restore proper transcription. A CHD8 protein that can bind but not remodel chromatin likely cannot mobilise nucleosomes, alter DNA accessibility, or grant transcription factors access to their sites, consistent with the established role of SNF2-family ATPases in nucleosome sliding, eviction, and histone-variant exchange [84]. The requirement for both domains - confirmed by a significant omnibus Friedman test and pairwise Wilcoxon comparisons across constructs (**Figure 7A**) - shows that CHD8 function depends on the coordinated action of H3K4me3 mediated recruitment and remodelling activities together. Unlike some other chromatin remodellers, in which recruitment and catalytic activity can substitute for one another [85], CHD8’s two modules appear non-interchangeable, possibly reflecting the precise temporal control needed during neuronal differentiation. The asymmetric distribution of CHD8 target genes across five domain-requirement categories - globally rescued, helicase-dependent, chromodomain-dependent, dual-domain-dependent, and low/no rescue (confirmed by chi-square test, **Figure 7B**) - further shows that different subsets of CHD8 targets have distinct structural requirements for transcriptional recovery. Median ATAC-seq log_2_FC at proximal peaks did not differ across these categories, consistent with our earlier observation that proximity to accessibility loss does not strongly predict rescue outcome (**Figure 7C**). To test whether these categories instead reflect differences in CHD8 occupancy or chromatin accessibility, we compared ChIP-seq binding intensity and ATAC-seq accessibility changes across all five groups among KO-dysregulated genes (n = 2,927; |log_2_FC| > 0.5). All five categories showed equivalent CHD8 occupancy - every gene in every group carried a CHD8 peak within 2 kb of its TSS - and comparable CHD8 peak scores (**Figure 7D**), indicating that domain-requirement category is independent of CHD8 occupancy or accessibility change.

### CHD8 regulates chromatin accessibility to control cell-fate decisions

Our ATAC-seq data show that CHD8 loss produces a chromatin accessibility landscape that is balanced in aggregate but skewed toward loss among high-confidence changes: across all 74,486 tested regions, global accessibility change was negligible (median LFC = +0.014, **Figure 6A**), WT and CHD8-KO NPCs remained strongly concordant genome-wide (Pearson r = 0.987; **Figure 6B**).However, among the subset of regions reaching statistical significance (padj < 0.05, |LFC| > 0.5; n = 4,486), accessibility loss clearly predominated (70.5% lost vs. 29.5% gained; **Figure 6C**). This indicates that CHD8 loss does not trigger wholesale chromatin remodelling but instead produces locus-specific, quantitative changes concentrated at particular regulatory elements. This pattern fits a model in which CHD8 acts predominantly to maintain accessibility at a specific, statistically robust subset of loci with a smaller counteracting set of loci at which it restrains accessibility, together channelling cells toward appropriate developmental fates.

Multi-omics integration illustrates just how locus-specific these effects are: only 8 genes were simultaneously CHD8-bound, transcriptionally dysregulated, and associated with an accessibility change in KO cells (**Supplementary Figure 1**). Although this convergence of three independent regulatory layers marks a very stringent, high-confidence set of direct CHD8 targets, it represents a small minority of total CHD8 binding events in NPCs - most CHD8-occupied loci show no transcriptional dysregulation upon CHD8 loss (**Figure 2C**), so CHD8 binding alone is not sufficient to confer transcriptional dependency. This fits prior observations that CHD8 occupancy is widespread but functionally selective, reflecting context-dependent requirements for remodelling activity that are not applied uniformly across all binding sites [14,17]. The broader overlap pattern - 246 genes shared between RNA-seq and ChIP-seq, 22 between RNA-seq and ATAC-seq, and 154 between ChIP-seq and ATAC-seq beyond the triple overlap - reflects the layered nature of CHD8’s regulatory influence, in which chromatin occupancy, accessibility remodelling, and transcriptional control are partially overlapping but mechanistically distinct. The 8-gene triple-overlap set should therefore be read as loci where all three regulatory outputs converge under stringent, statistically supported thresholds, not as the totality of CHD8’s regulatory activity. This picture is consistent with previous reports that CHD8 recruits histone H1 to repress beta-catenin-dependent transcription [86], supporting a model in which CHD8 coordinates multiple chromatin-regulatory mechanisms to steer cell-fate decisions during neuronal differentiation.

## Conclusions

Through comprehensive multi-omics profiling of female mouse ES cells undergoing neuronal differentiation, we have revealed that CHD8 functions as a key regulator of female neuronal differentiation programs. CHD8 undergoes extensive cistrome remodelling during the ES-to-NPC transition, with the majority of NPC binding sites localising to distal regulatory elements alongside a substantial fraction at H3K4me3-marked active promoters, where it regulates expression of genes critical for neuronal differentiation, synaptogenesis, and ASD-relevant pathways. Both CHD8’s chromodomains (for H3K4me3-dependent recruitment) and helicase domain (for chromatin remodelling) make distinct and non-redundant contributions to transcriptional rescue of direct NPC target genes, with different subsets of CHD8 targets showing separable domain requirements for effective transcriptional recovery.

CHD8 loss causes widespread transcriptional dysregulation affecting female NPCs, with a consistent downregulation-predominant asymmetry in the knockout line and the stronger knockdown line, and pathway enrichment analysis revealing broad suppression of ribosomal genes expression, ion transport, and neurodevelopmental programmes among downregulated genes. The magnitude of dysregulation across these perturbation conditions highlights the importance of including female-derived systems in neurodevelopmental disorder research. Integration of CHD8 binding, gene expression, and chromatin accessibility data identifies 8 high-confidence genes simultaneously CHD8-bound, transcriptionally dysregulated, and associated with significant chromatin accessibility changes in KO cells, providing high-confidence candidate loci at which CHD8 occupancy, chromatin accessibility and transcriptional regulation converge.

## Abbreviations

ASD: Autism spectrum disorder
ATAC-seq: Assay for Transposase-Accessible Chromatin with high-throughput sequencing
CHD8: Chromodomain Helicase DNA-binding protein 8
ES: Embryonic stem cells
NPC: Neural progenitor cells
XCI: X-chromosome inactivation
Xist: X-inactive specific transcript
KO: Knockout
KD: Knockdown
FL: Full-length
ΔChromo: Chromodomain deletion mutant
ΔHelicase: Helicase domain deletion mutant
ChIP-seq: Chromatin immunoprecipitation sequencing
RNA-seq: RNA sequencing
DEG: Differentially expressed genes
GO: Gene ontology
GSEA: Gene set enrichment analysis
DA: Differentially accessible
TSS: Transcription start site
PCA: Principal component analysis
FDR: False discovery rate
SFARI: Simons Foundation Autism Research Initiative

## Declarations

### Ethics approval and consent to participate

Not applicable (study used established mouse ES cell lines only).

### Consent for publication

Not applicable.

### Availability of data and materials

All high-throughput sequencing data generated in this study have been deposited in the Gene Expression Omnibus (GEO) database under accession numbers GSE166858 and GSE166859 (previously published datasets from Cerase et al., 2021). The new RNA-seq datasets from rescue experiments have been deposited in GEO under accession number GSE346086. All data supporting the conclusions of this study are included within the article and its supplementary files. Code used for the analysis has been deposited in GitHub (https://github.com/ajaykumardanga-lgtm/chd8-chip-atac-rnaseq-pipeline).

### Competing interests

The authors declare no competing interests.

### Funding

This work was funded by EMBL recurrent funding, QMUL intramural support, Barts Charity grants, Rett Syndrome Research Trust and Loulou Foundation and University of Pisa intramural funding (to A.C.) and A.K.D. was supported by a Department of Biotechnology - Research Associateship (DBT-RA).

### Authors’ contributions

M.M.F contributed to manuscript writing and some analysis. A.K.D and A.Co. performed all bioinformatic analyses. A.Co., N.P. and M.M.F. verified the correctness of all the bioinfomatic analysis. F.E. contributed to the statistical and methodological evaluation of the study, assessed the internal consistency of the reported analyses, and critically reviewed and revised the manuscript. G.G.T and A.Ce. conceived and supervised the study, A.C. performed the experimental work, generated cell lines and rescue constructs, interpreted results, and wrote the manuscript with A.K.D. All authors read and approved the final manuscript.

## Acknowledgements

We thank Marta Biagioli for critical reading of our manuscript. Philip Avner for support and guidance, Alexander N. Young and Nerea Blanes Ruiz for generating and characterizing cell lines used in this study, Andreas Buness for initial bioinformatic analyses, and all members of the Cerase laboratory for discussions. We acknowledge Queen Mary University of London Genome Centre for sequencing services and High-Performance Computing facilities for computational resources.

## Notes

https://www.ncbi.nlm.nih.gov/geo/query/acc.cgi?acc=GSE346086

https://github.com/ajaykumardanga-lgtm/chd8-chip-atac-rnaseq-pipeline

### Competing Interest Statement

The authors have declared no competing interest.

